# Species Delimitation in Indian *Ledebouria*: Resolving the putative species status of infra-generic taxon Despite Phylogenomic Discordance

**DOI:** 10.64898/2026.06.18.733124

**Authors:** Sachidanand Nayak, Pradip Deshmukh, Shrirang R. Yadav, Manoj M. Lekhak, Siddharthan Surveswaran

## Abstract

*Ledebouria* is a genus of geophytic herbs distributed mainly in Africa, with a few species also occurring in the Arabian Peninsula, Madagascar, India, and Sri Lanka. For many years, South Asian accessions were identified as *Ledebouria revoluta* until recent molecular studies demonstrated that the African *L*. *revoluta* is distinct from the taxa occurring in South Asia, leading to the resurrection of the name *L*. *hyacinthina*. Despite this taxonomic revision, the molecular and morphological relationships among the diverse morphotypes and cytotypes within *L*. *hyacinthina* remain poorly resolved. Because plastid DNA markers have shown limited ability to detect variation within this group, we employed the Angiosperms353 target enrichment marker set, comprising hundreds of low-copy nuclear genes, to investigate phylogenetic relationships among eight representative accessions. These were analysed together with published global datasets generated using the same marker set. Our results support the reinstatement of *L. hyacinthina var. hyderabadensis* to the status of a species., whereas the remaining accessions analysed showed very high phylogenomic discordance. Overall, our findings suggest that molecular divergence within this species complex remains limited, with both the Angiosperms353 nuclear markers and chloroplast sequences providing insufficient resolution to fully under their evolutionary relationship.

## 1. Introduction

The genus *Ledebouria* Roth (Family: Asparagaceae *sensu* APG III (The Angiosperm Phylogeny Group, 2009), sub-family: Scilloideae and tribe: Hyacintheae) (Manning, 2020) consists of bulbous herbs, distributed mainly in the Sub-Saharan Africa with a maximum diversity in South Africa (POWO, 2026). However, a few species occur outside continental Africa, including in the Arabian peninsula, Socotra, Madagascar and peninsular India (Howard et al., 2022; POWO, 2026). The only widespread species was considered to be *Ledebouria revoluta* (L.f.) Jessop which is distributed from Eritrea to S. Africa, SW. Arabian Peninsula, India and Sri Lanka, however the understanding of this species has been changed with recent work (Howard et al., 2022).

*Ledebouria* was first described by Roth (Roth, 1821) from India with *Ledebouria hyacinthina* Roth as the type species. *Ledebouria* was then moved to the genus *Scilla* L. by Baker (Baker, 1872). Following this, the name *Scilla* was used to describe species under this group. Later, Baker (Baker, 1896) treated a majority of the species formerly treated under *Scilla* under subgenus *Ledebouria* to differentiate it from subgenus *Euscilla*. Based on the study of the plant bulbs with the deciduous leaves, the structure of the ovary, particularly, the placement of the ovules in each locule, Jessop (Jessop, 1970) re-established the group as its own genus. This was further proved by cladistic analysis of morphological data as well as plastid introns and intergenic marker sequence data (Stedje, 1998).

*Drimiopsis*, *Ledebouria* and *Resnova* are the most closely related genera among Massonieae and are treated as separate genera (Lebatha et al., 2006) or treated as sections under *Ledebouria* (Buerki et al., 2012; Manning and Goldblatt, 2012; Manning, 2020; Manning et al., 2003; Pfosser et al., 2003). Currently, POWO recognizes 64species of *Ledebouria*, nine species of *Drimiopsis* and three species of *Resnova* (POWO, 2026).

Howard et al. (Howard et al., 2022) recognized the difficulty in reconstructing a robust evolutionary understanding of the three genera due to homoplasies and a very limited number of molecular markers, particularly plastid markers which could not trace genomic recombinations. Therefore, they used a target enrichment sequencing of nuclear markers using the Angiosperms353 marker set (Johnson et al., 2019). Their work revealed four clades in Ledebouriineae, two in *Ledebouria* and one each in *Drimiopsis* and *Resnova*. Among them, one *Ledebouria* clade was entirely sub-Saharan African and the other was East-African and non-African. Their work also showed that Indian and Sri Lankan *Ledebouria revoluta* was in the latter clade suggesting the African and Asian *L. revoluta* are different entities.

Prior to Howard et al., (2022), Giranje and Nandikar (Giranje and Nandikar, 2016) recognised four species in India, *L. viridis*, *L. revoluta*, *L. karnatakensis* and *L. hyderabadensis* based on morphological characters. Chakral et al. (Chakral et al., 2024) used plastid molecular data to distinguish African *L*. *revoluta* from Indian *L*. *revoluta* and thereby resurrected the species name from *Scilla hyacinthina*, which had been synonymized (Ramana et al., 2012; Stedje and Thulin, 1995) with *L. revoluta* (L. f.) Jessop since Jessop (Jessop, 1970). Their analysis echoed the findings of Howard et al. (2022) in distinguishing *L*. *revoluta* from the Indian entity; however, the latter was not cited. Chakral et al. (2024) also treated the former species *L*. *hyderabadensis* and *L*. *karnatakensis* as a variety and a forma of *L*. *hyacinthina*, respectively. Moreover, they described one new variety and one new forma, *L*. *hyacinthina* var. *obtusata* and *L*. *hyacinthina* f. *recurvata*, respectively. Their work was based on morphological analysis as well as the analysis of two plastid intergenic regions, *trnL*–*trnF* and *trnC*–*ycf6*.

Previous research has identified ten distinct cytotypes for *L. hyacinthina* (current name for Indian specimens of *L. revoluta)*, with reported chromosome counts (2*n*) including 28, 30, 44, 45, 46, 58, 60, 64, 68, and 70 (Dixit et al., 1989). A first record of hexaploidy (2n = 90) in *L*. *revoluta* (currently treated as *L*. *hyacinthina*) has been reported, along with a highly asymmetrical (2C) karyotype (Giranje et al., 2019). Diploid populations (2n = 30) are typically found in plain regions and primarily exhibit sexual reproduction. Triploid (2n = 45) and hexaploid (2n = 90) populations are so far known only from higher altitudes in the Western Ghats. These populations may be adapted to high-rainfall zones and predominantly reproduce vegetatively through leaf-tip bulbils (Giranje et al., 2019). Given the different cytotypes and morphological ambiguity, we aimed to test the species delimitation of Indian *Ledebouria* to determine whether distinct karyotypes and morphotypes could be resolved using molecular phylogenetics. For this, we used low copy nuclear markers from the Angiosperms353 probe set. Furthermore, we sought to compare whether the results were congruent with plastid markers, *trnL* intron and *trnL-trnF* and partial *matK*.

## 2. Materials and Methods

### 2.1 Sample collection and target sequencing

*Ledebouria* samples were collected from various locations in peninsular India and grown in the botanical garden at Shivaji University, Kolhapur, India. All samples were deposited in the herbarium of Shivaji University (SUK) (Table 1). Among them, eight samples representing different cytotypes were used for target capture sequencing. DNA extraction, library preparation, target-capture and NGS sequencing was performed by Nucleome Informatics, Hyderabad, India (www.nucleomeinfo.com). DNA was isolated from silica gel dried leaf tissues using the CTAB method. The quantification and purity ratios of extracted DNA were checked using NanoDrop 2000 Spectrophotometer (Thermo Fisher Scientific, MA, USA). The integrity of the DNA was checked using Agarose Gel Electrophoresis (1% Agarose Gel). Library Preparation was done using the Kapa Hyperplus Kit (Roche, USA) and library purification was done using Kapa Pure Beads (Roche, USA). Library length was assessed using the DNA High-sensitivity chip using the Agilent 2100 Bio-analyzer (Agilent, USA). The prepared library was multiplexed and hybridized with the myBaits kits expert kit (Daicel Arbor Biosciences, USA) according to the manufacturer’s instructions. Hybridization was carried out at 65°C for 24 hours. The target captured library was amplified and sequenced using the NovaSeq 6000 (Illumina, USA).

**Table 1.**
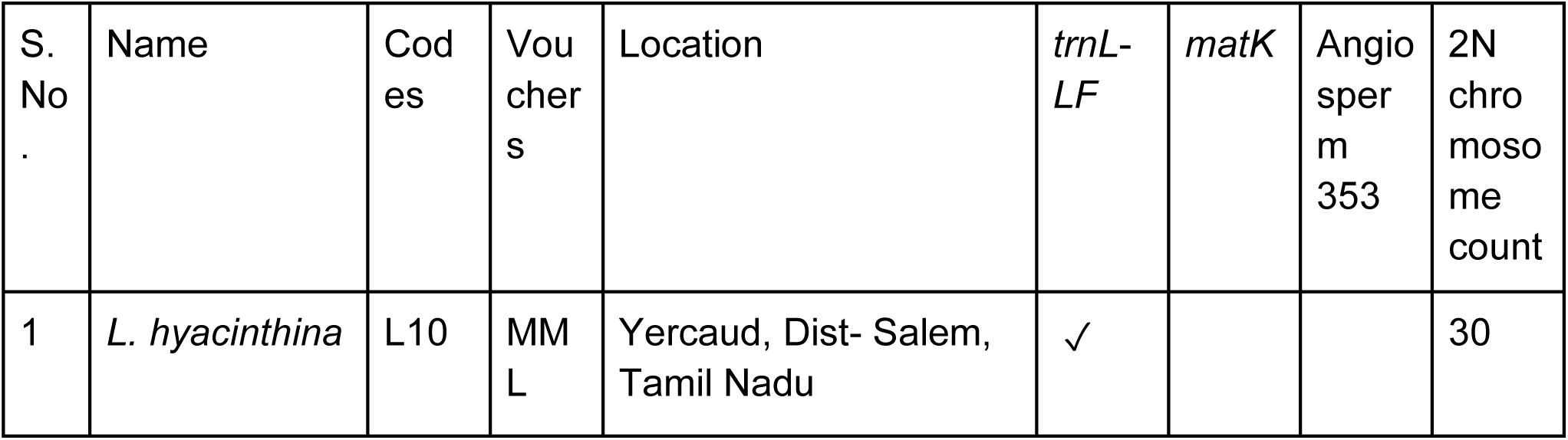
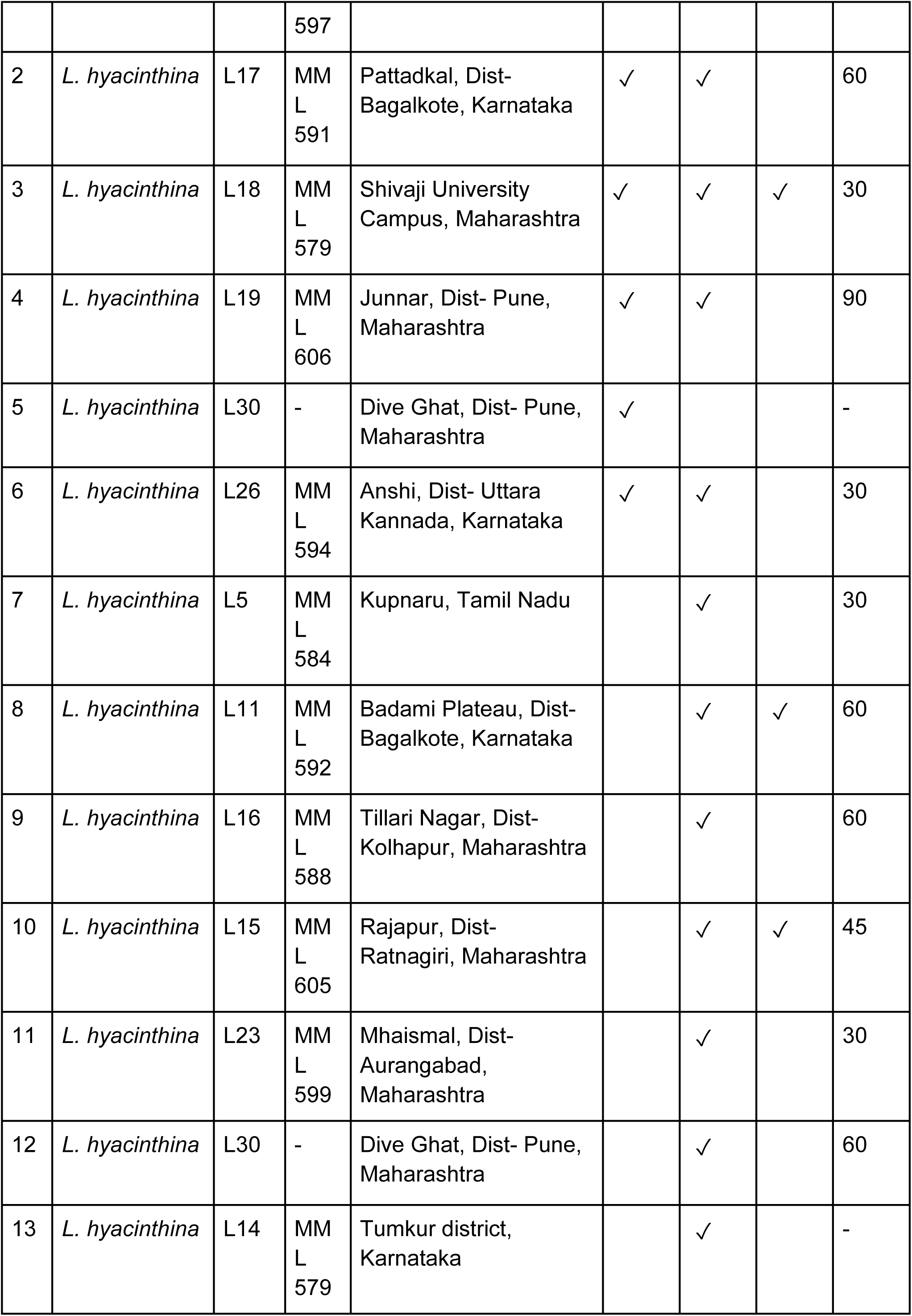
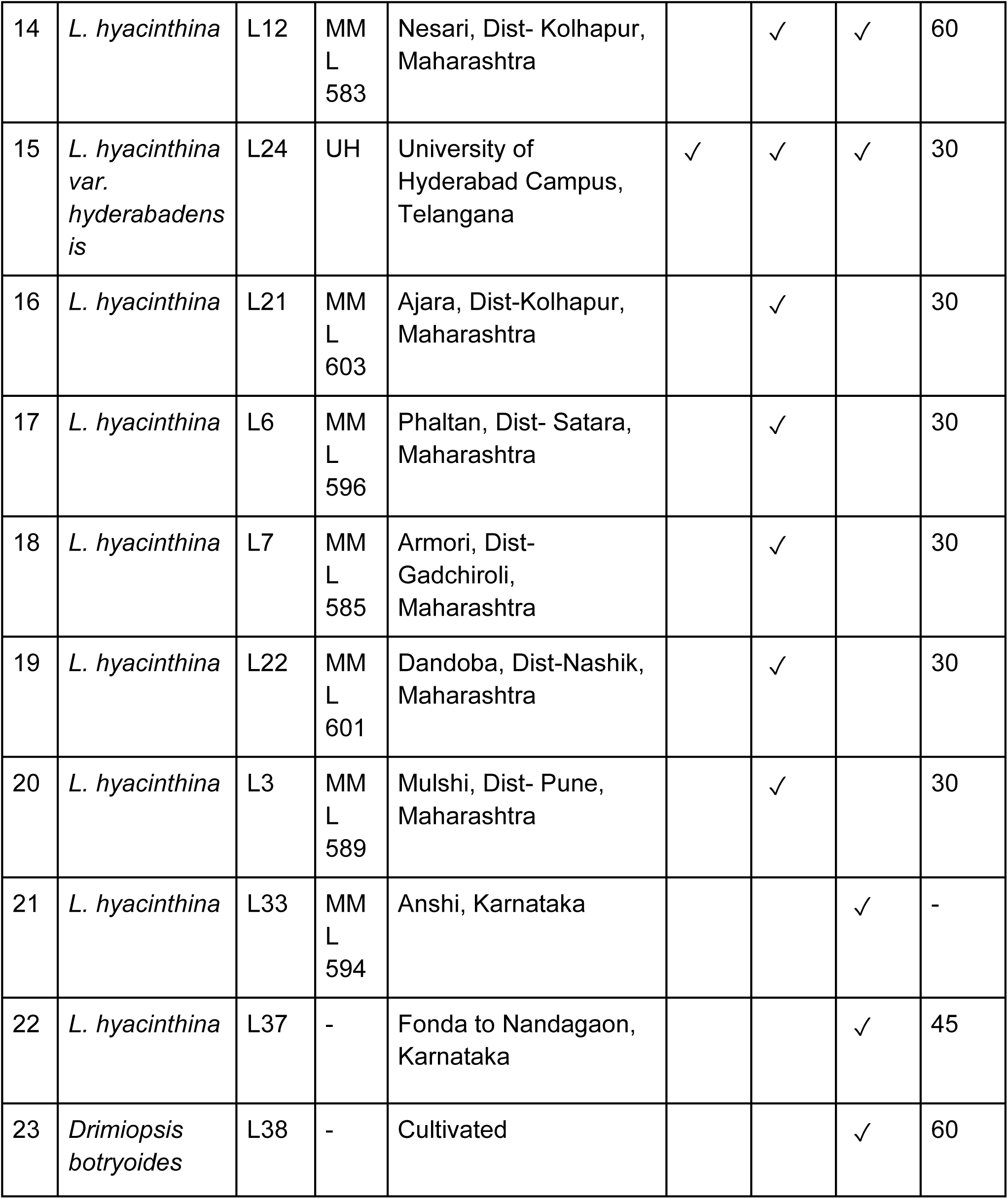
Samples used for Angiosperm353 and/or *matK* and *trnL*-*LF* analysis. Chromosome counts are from Deshmukh (2023).

### 2.2 Sequence pre-processing, assembly, and alignment

Paired-end raw reads of the 8 plant samples were trimmed using Trimmomatic-0.39 (Bolger et al. 2014) to remove adapters and low-quality reads (*LEADING:20 TRAILING:20 SLIDINGWINDOW:4:20 MINLEN:50*). The read pairs where both reads passed the described filtering parameters were retained.

These high-quality paired-end reads were then assembled by HybPiper version 2.3.2 (Johnson et al., 2016) (subcommand: *assemble*) using BWA (Li, 2013) to map the reads to the Angiosperms353 target sequences (Johnson et al., 2019). The maximum target sequence to be saved in BLASTx search was set to 50. From the resulting assembly, supercontigs (sequences containing both exons and introns) were retrieved by HybPiper version 2.3.2 (subcommand: *retrieve_sequences*), excluding the putative chimeric genes. Additionally, the potential paralogs were identified by HybPiper version 2.3.2 (subcommand: *paralog_retriever*).

The supercontig supermatrix and its corresponding partition file for the TAXA70 dataset (the plant samples with less than 70% gaps/ambiguities in the supermatrix) were downloaded from Howard et al. (2022). The supercontig supermatrix was subsequently split into individual gene alignments based on the partition file. The supercontigs retrieved from our 8 samples were then aligned to their corresponding gene alignments of the TAXA70 dataset, preserving the original alignment structure. This was performed using MAFFT v7.525 (Katoh et al., 2019) with the parameters *--add, --auto, --ep 0.123* and *--op 2*. For any of the genes in our 8 samples that could not be retrieved or exhibited paralogy issues, the corresponding alignment positions were filled with “N” characters in the respective gene alignment files to account for missing data during gene and species tree reconstruction.

### 2.3 Gene and species tree reconstruction

Following data cleaning, two datasets were analysed: (1) a full dataset of 334 genes, representing maximal gene recovery across all samples (including those from Howard et al., 2022), named “untrimmed full dataset” and (2) a reduced dataset of 106 genes, corresponding to the maximum gene recovery observed in *L*. *hyacinthina* var. *hyderabadensis* named “untrimmed reduced dataset”, Both datasets were trimmed using TrimAl v1.5 (Capella-Gutiérrez et al., 2009) with the automated method (-automated1), retaining sequences even if they were composed entirely of gaps (-keepseqs). The resulting datasets were called (3) “trimmed full dataset” and (4) “trimmed reduced dataset”. Thus four datasets were used for further phylogenetic analyses.

Gene trees for each dataset were inferred using IQ-TREE version 3.0.1 (Wong et al., 2025). The ultrafast bootstrap approximation (UFBoot2) (Hoang et al., 2018) was implemented with 1000 replicates along with ModelFinder Plus (Kalyaanamoorthy et al., 2017) for model selection (*-B 1000 -m MFP*). An edge-linked proportional partition model (Chernomor et al. 2016) was applied, with separate tree inference for each partition (*-S*).

For each dataset, species trees were independently constructed from the corresponding gene trees or gene alignments using IQ-TREE version 3.0.1, ASTRAL-IV v1.23.4.6 (which uses quartet-based species tree inference method) (Zhang et al., 2025) (Tabatabaee et al., 2023) Weighted ASTRAL v1.23.3.7 (which uses threshold-free weighting schemes for the quartet-based species tree inference to reduce the impact of quartets with low support or long terminal branches or both) (Zhang and Mirarab, 2022), and Asteroid v1.1 (Accurate Species Tree Estimation RObust to Incomplete Data sampling) (Morel et al., 2023) resulting in four alternative species tree topologies per dataset. The IQ-TREE version 3.0.1 was run with SH approximate likelihood ratio tests (SH-aLRT) and ultrafast bootstrap (uBS) values. (Guindon et al., 2010), each with 1000 replicates for calculating branch support (*-B 1000 --alrt 1000*). Moreover, to identify the optimal partitioning scheme, *TESTMERGE* (Lanfear et al., 2012) was implemented with the relaxed hierarchical clustering algorithm (Lanfear et al., 2014), constraining the search to top 10% of partition merging schemes to reduce the computational burden (*-m TESTMERGE --rcluster 10*). Additionally, both the partitions and the sites within the resampled partitions were resampled during the bootstrap analysis (Seo et al., 2005); (Gadagkar et al., 2005) (*--sampling GENESITE*). ASTRAL-IV v1.23.4.6 and Weighted ASTRAL v1.23.3.7 were executed with default parameters. Node support, local posterior probabilities (LPP) were generated in ASTRAL and Weighted ASTRAL. Asteroid v1.1 was executed using gene tree branch lengths to build distance matrices (--use-gene-bl) and by computing 1000 bootstrap trees (-b 1000).

### 2.4 Concordance analysis

The gene concordance factor (gCF) and site concordance factor (sCF) (Minh et al., 2020) were quantified using IQ-TREE version 3.0.1 for all the datasets separately. The species tree reconstructed by IQ-TREE version 3.0.1 was used as the reference tree to which gCF and sCF values were assigned. The gene trees, also generated by IQ-TREE version 3.0.1, were set as source trees for computing gCF. For sCF estimation, 100 quartets were randomly sampled per internal branch (*--scfl 100*).

### 2.5 Species delimitation and migration rate analysis

Species delimitation analysis was conducted using Bayesian Phylogenetics and Phylogeography (BPP) v4.8.7 (Flouri et al., 2018) under the Multispecies Coalescent (MSC) model. An A10 analysis (species delimitation using a fixed guide tree) was conducted using a subset of trimmed reduced dataset (106 loci) comprising of the *Ledebouria* samples used in this study, *Ledebouria revoluta* samples from India and Sri Lanka as well as Ledebouria_kirkii_CCH159_Tanzania and Ledebouria_sp_CCH161_Tanzania collected by Howard et al. (2022) by rjMCMC (reversible-jump Markov chain Monte Carlo) algorithm 1 to test species boundary among five putative species (based on phylogenetic analysis); i) *Ledebouria revoluta*: *Ledebouria revoluta* samples from India and Sri Lanka collected by Howard et al. (2022) (3 samples), ii) *Ledebouria hyacinthina*: *Ledebouria* samples collected in this study except *Ledebouria hyacinthina var. hyderabadensis* (6 samples), iii) *Ledebouria hyderabadensis*: *Ledebouria hyacinthina var. hyderabadensis* (1 sample), iv) *Ledebouria kirkii:* Ledebouria_kirkii_CCH159_Tanzania (1 sample) and v) *Ledebouria sp.*: Ledebouria_sp CCH161_Tanzania (1 sample) using the fixed guide tree: *(((Ledebouria hyacinthina, Ledebouria revoluta), Ledebouria hyderabenesis), (Ledebouria kirkii, Ledebouria sp));*. Three different *thetaprior* and *tauprior* parameter settings were used for species delimitation to check sensitivity to the priors (see Data S1 for control and other input files used for species delimitation analysis). Each parameter setting was run twice using different seed values to check reproducibility.

Similarly, migration rate estimation was performed using BPP v4.8.7 under MSC model with migration (MSC-M) (Flouri et al., 2023) with four different settings to check sensitivity to the priors (see Data S1 for control and other input files used for gene flow estimation). Each parameter setting was run twice using different seed values to check reproducibility.

### 2.6 Plastid DNA sequence amplification and sequencing

The plastid genes, partial *matK*, *trnL* intron and *trnL-trnF* intergenic spacer (hereafter called as *trnL-LF*) were PCR amplified using primers XF, 5R (*matK*) and cf (*trnL-LF*) (Cuénoud et al., 2002; Ford et al., 2009; Taberlet et al., 1991). PCRs in 25 μL reactions with ∼20 ng DNA, 40 amplification cycles, with an annealing temperature of 58 °C. PCR products were checked on by agarose gel electrophoresis. PCR product purification and Sanger sequencing were performed by Barcode Biosciences Pvt. Ltd (https://www.barcodebiosciences.com/). Bidirectional reads were quality-checked and assembled into consensus contigs using the Staden package (Staden et al., 2000). Sequences were aligned with MUSCLE in AliView (Larsson, 2014) followed by manual refinement. Substitution models selected under the Akaike Information Criterion (AIC) using jModelTest 2 (Darriba et al., 2012). Bayesian phylogenetic inference was performed in MrBayes 3.2.7a (Ronquist et al., 2012) with two independent runs of 2 million generations, sampling every 1,000 generations. The convergence of the runs were checked using Tracer 1.7.2 (Rambaut et al., 2018) and the first 10 % trees were removed as burn-in. Trees were visualized using FigTree 1.4.4 (Rambaut, 2018) and edited for publication using the InkScape 1.4.3 (inkscape.org) vector editing program.

## 3. Results

### 3.1 Phylogenetic placement of Indian Ledebouria within Ledebouriinae

In order to understand the phylogenetic relationship and species boundaries of the Indian *Ledebouria* taxon with other members of *Ledebouriinae,* 8 samples collected from various locations in peninsular India representing different cytotypes were used for Angiosperms353 target capture sequencing (Table 1). In addition to these 8 samples, 76 samples were incorporated from Howard et al. (2022). We have used the name *Ledebouria hyacinthina* for the Indian representatives of *Ledebouria sensu* Chakral et al. (2024) but retained the name *L. revoluta* for Indian and Sri Lankan accessions obtained from Howard et al. (2022).

In all the samples generated in this study >70% of the reads were mapped to target sequences and <7 target sequences were flagged for paralogy in each sample (Table S1). The sample, namely *L. hyacinthina* var. *hyderabadensis* provided less yield (106 supercontigs and 31147 bases recovered) than other samples (322-338 supercontigs and 275821-639365 bases recovered) (TableS1).

In order to reduce the impact of *L. hyacinthina* var. *hyderabadensis* sample’s low yield on downstream analyses, two additional datasets were generated namely, i) Untrimmed reduced dataset: a reduced dataset of 106 genes, corresponding to the maximum gene recovery observed in *L*. *hyacinthina* var. *hyderabadensis* and ii) Trimmed reduced dataset: obtained after trimming the untrimmed reduced dataset, in addition to iii) Untrimmed full dataset: a full dataset of 334 genes, representing maximal gene recovery across all samples (including those from Howard et al., 2022) and iv) Trimmed full dataset: obtained after trimming the untrimmed full dataset (see methods for details).

The species trees inferred from each dataset described earlier, using IQ-TREE and ASTRAL-IV consistently placed Indian *L. hyacinthina* within *Ledebouria* clade B with strong branch support values (Fig. 1-2; Fig. S1-S6). The *Drimiopsis botryoides* sample collected from India in this study was nested within the corresponding *D. botryoides* clade of Howard et al. (2022) as expected (Fig. 1-2; Fig. S1-S6). We have designated this well-supported lineage of *Ledebouria* consisting of 7 samples from India collected in this study as well as *Ledebouria revoluta* samples from India and Sri Lanka collected by Howard et al. (2022) as the South Asian clade (SA clade)(Fig. 1-2; Fig. S1-S6).

**Fig. 1.**
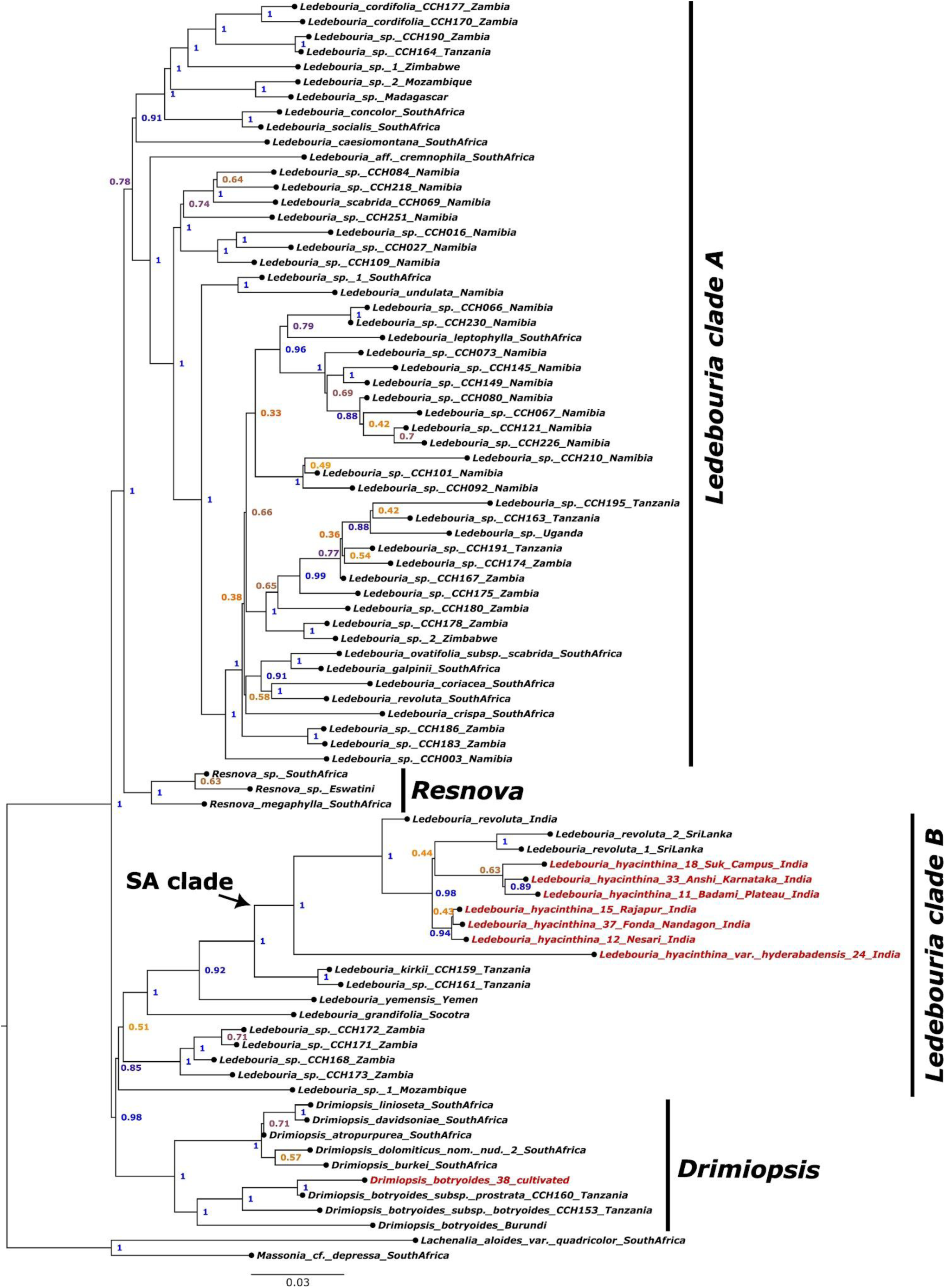
Species tree of *Ledebouria* taxa generated by ASTRAL using the trimmed full dataset containing 334 genes. Numbers at nodes indicate local posterior probability (LPP) support. Sequences generated for this study are indicated in red.

**Fig. 2.**
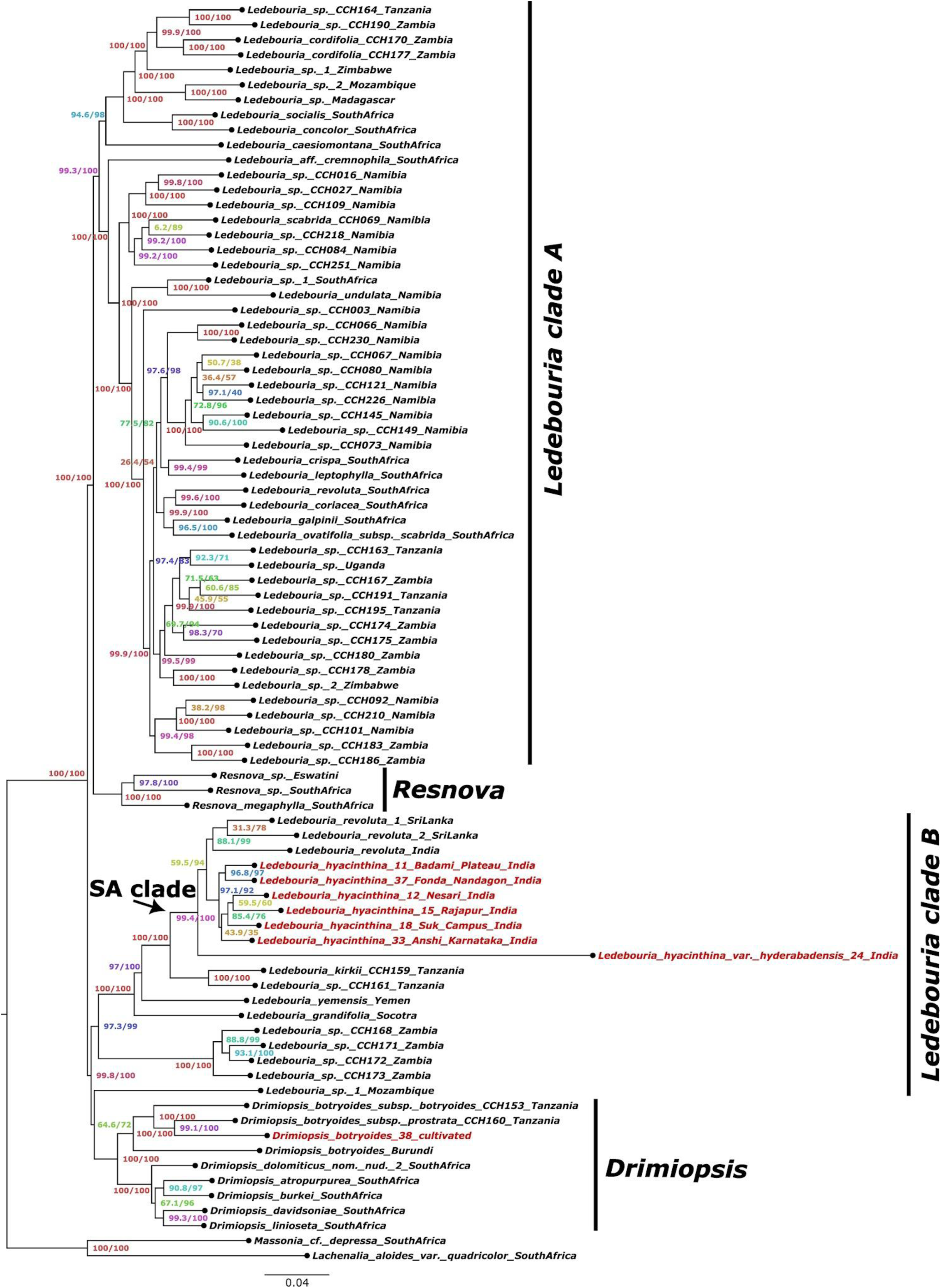
Species tree of *Ledebouria* taxa generated by IQTREE using the trimmed full dataset containing 334 genes. Numbers at nodes indicate SH-aLRT (SH approximate likelihood ratio tests) and ultrafast bootstrap values. Sequences generated for this study are indicated in red.

### 3.2 Multiple evidence for the recognition of L. hyacinthina var. hyderabadensis as a putative species-level taxon

Given the morphological distinction of *L. hyacinthina* var. *hyderabadensis* among Indian accessions, characterized by fewer than 25 flowers per inflorescence (compared to at least 40 in others), presence of sterile flowers, and a beaked ovary (Chakral et al. 2024); we evaluated its taxonomic status using the molecular evidence derived from the Angiosperms353 target capture sequencing to determine whether it represents a species-level taxon.

Because the sample of *L. hyacinthina* var. *hyderabadensis* was of comparatively low yield, two additional datasets (untrimmed and trimmed reduced datasets) were also generated and analysed as described in detail in the previous section and methodology. In the species trees inferred by IQ-TREE and ASTRAL-IV using the trimmed and untrimmed dataset of both 106 and 334 genes showed that *L. hyacinthina* var. *hyderabadensis* has a slightly elongated branch and occupied an early-diverging position within the SA clade with very high branch support values (Fig. 1-2; Fig. S1-S6). To determine if the missing data, low support and long terminal branches in gene trees due to *L. hyacinthina* var. *hyderabadensis* sample, was responsible for elongated branch and early-divergence in the species tree, Asteroid (an algorithm specifically designed to robustly infer species tree from gene trees under high proportions of missing data) and Weighted ASTRAL (which uses threshold-free weighting schemes for the quartet-based species tree inference to reduce the impact of quartets with low support or long terminal branches or both) were additionally used on all four datasets for species tree inference. The species trees inferred by Weighted ASTRAL still showed the early divergence of *L. hyacinthina* var. *hyderabadensis* within SA clade, though branch length had reduced comparatively (Fig. 3; Fig. S7-S9). Out of 4 species trees generated by Asteroid using the four datasets, two showed early divergence of *L. hyacinthina* var. *hyderabadensis* within SA clade with comparatively reduced branch length and good support values (Fig. S10-S11). The other two also showed *L. hyacinthina* var. *hyderabadensis* diverged before other members of SA clade but their support values were poor (Fig. S12-13) and in one of them the placement of *L. hyacinthina* var. *hyderabadensis* differed (Fig. S13). Furthermore, the overall topology of the species trees inferred using Asteroid (Fig. S10-S13) were different from the ones inferred by IQ-TREE, ASTRAL-IV and Weighted ASTRAL, which showed consistent topologies (Fig. 1-3; Fig. S1-S9).

**Fig. 3.**
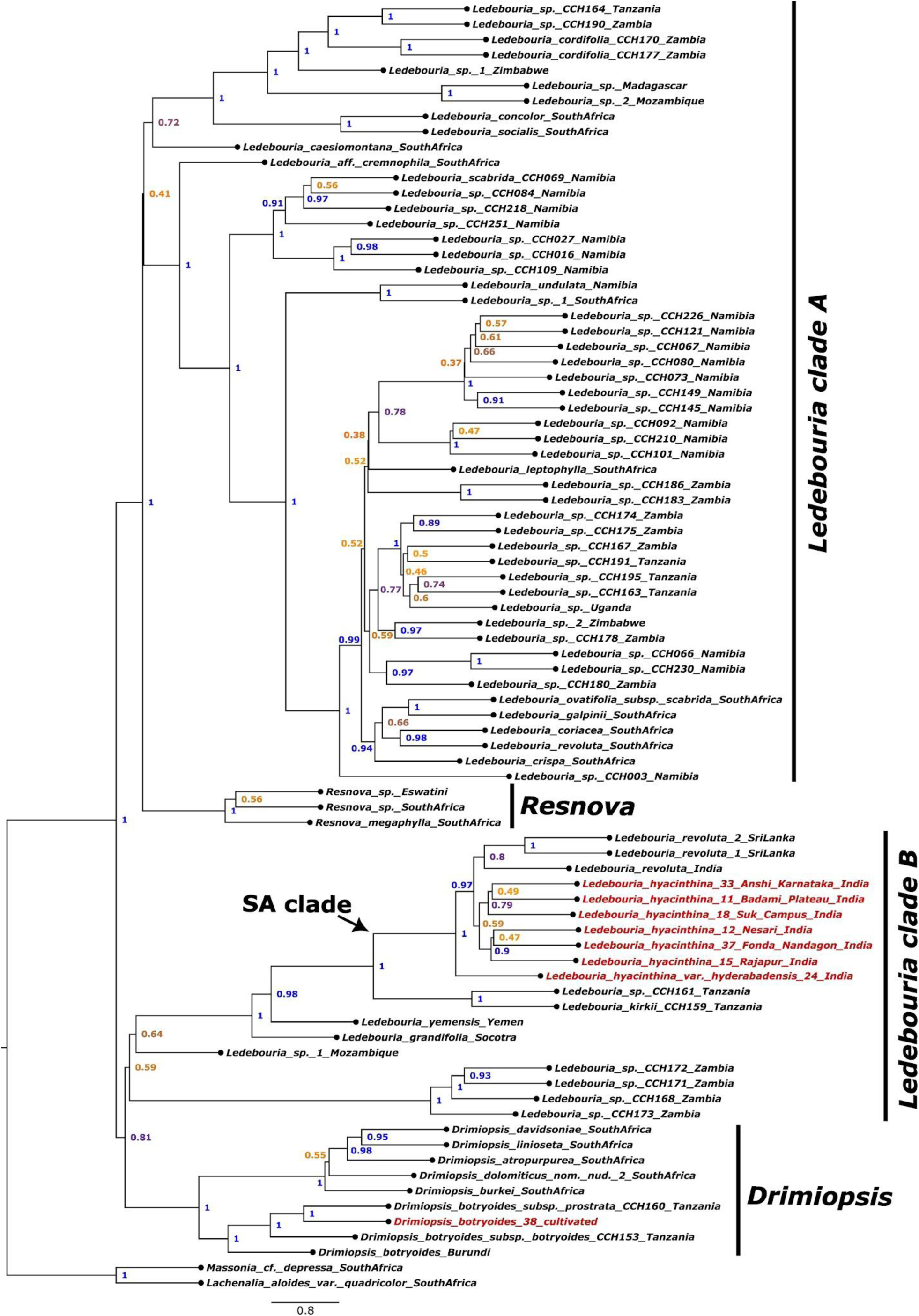
Weighted ASTRAL generated species tree of *Ledebouria* taxa using a trimmed reduced dataset containing 106 genes. Numbers at nodes indicate local posterior probability (LPP) support. Sequences generated for this study are indicated in red.

Concordance analysis evaluates how much genomic data (gene trees and genomic sites) support a particular phylogenetic topology compared with alternative topology such as Nearest Neighbour Interchange topology 1 (NNI-1) and Nearest Neighbour Interchange topology 1 (NNI-2). For the four dataset used in this study, the NNI-1 and NNI-2 also supported early divergence of *L. hyacinthina* var. *hyderabadensis* from other members of SA clade (Fig. S14-S15). Based on the gene concordance factor (gCF) and site concordance factor (sCF) analysis of all four datasets (see methods), it was observed that ∼44-50% of the gene trees and ∼32-43% of sites support that *L. hyacinthina* var. *hyderabadensis* is diverging early in the SA clade (Fig. 4A-B; Fig. S16-S21); ∼22-26% gene trees and ∼20-25% sites support that *L. hyacinthina* var. *hyderabadensis* is diverging from the common ancestor of SA clade, *Ledebouria kirkii* (CCH159, Tanzania) and *Ledebouria sp.* (CCH159, Tanzania) (NNI-1) (Fig. 4A-B; Fig. S14, S16-S21); ∼0-3% gene trees and ∼35-47% sites support that *L. hyacinthina* var. *hyderabadensis* diverged early from the SA clade and recently diverging from the common ancestor of *Ledebouria kirkii* (CCH159,Tanzania) and *Ledebouria sp.* (CCH159,Tanzania) (NNI-2) (Fig. 4A-B; Fig. S15, S16-S21; Fig. S17), and ∼25-29% gene trees support polyphyly of *L. hyacinthina* var. *hyderabadensis* (Fig. 4A; Fig. S16-S18). The sCF analysis does not show values for polyphyly.

**Fig. 4.**
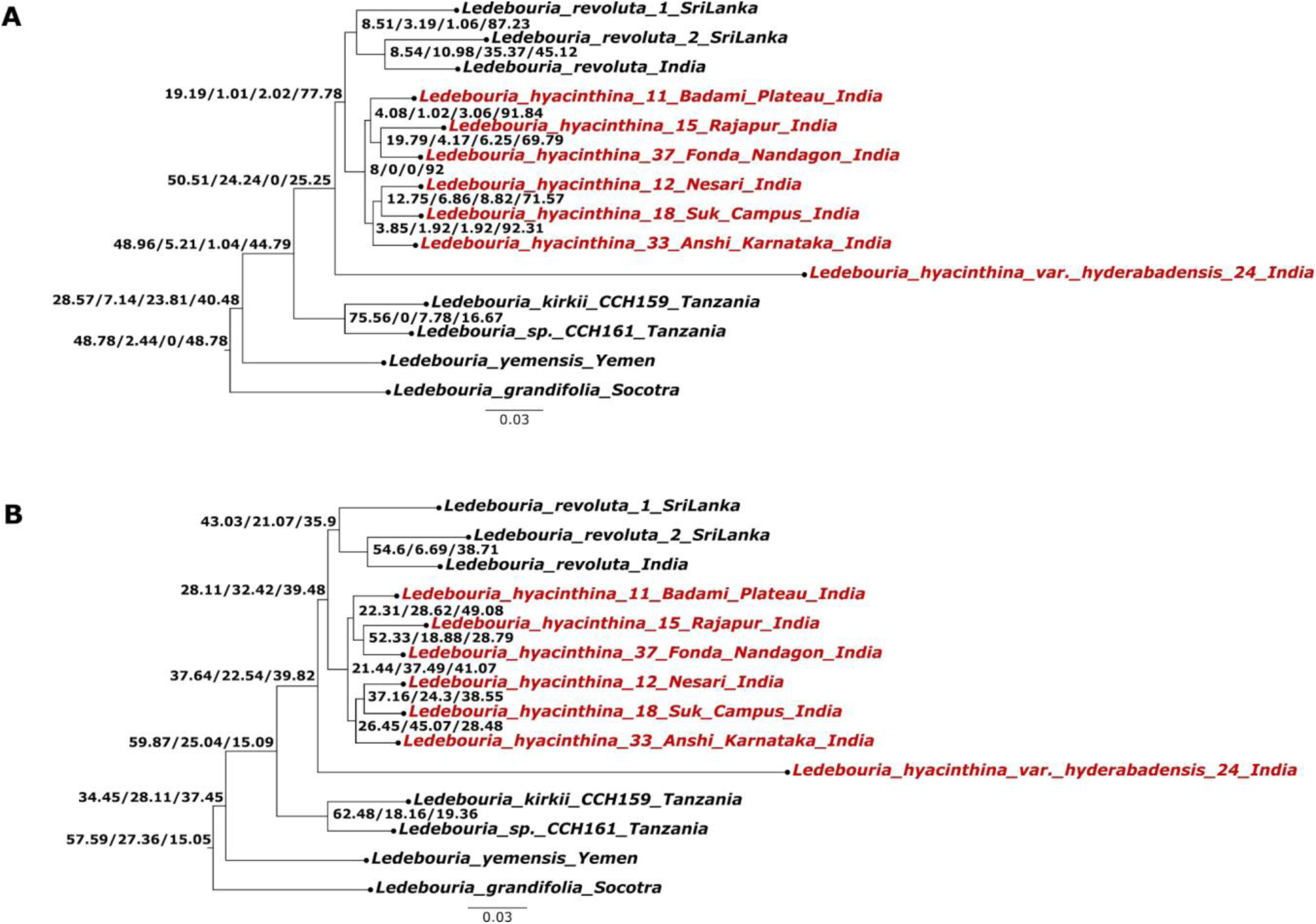
A. Gene concordance factor analysis of the *Ledebouria* SA clade using the untrimmed reduced dataset containing 106 genes. Numbers on the nodes indicate gene concordance factor/gene discordance factor for Nearest Neighbour Interchange (NNI)-1 branch/gene discordance factor for NNI-2 branch/gene discordance factor due to polyphyly. B. Site concordance factor analysis of the *Ledebouria* SA clade using the untrimmed reduced dataset containing 106 genes. Numbers on the nodes indicate site concordance factor averaged over 100 quartets/site discordance factor for alternative quartet 1/site discordance factor for alternative quartet 2. Sequences generated for this study are indicated in red.

The species delimitation analysis evaluated 7 alternative species-delimitation models under three different *thetaprior* and *tauprior* parameter settings, each repeated twice with different seed values (Table S2). In all six runs, the three selected putative species (based on inferred species tree) namely, *Ledebouria hyacinthina*, *Ledebouria revoluta* and *Ledebouria hyderabadensis* were consistently observed to be distinct species with posterior probability of 1 (Table S2; Fig. S22-S23).

Additionally, migration rate analysis was performed to check the gene flow between the selected putative species. Across all 8 independent runs, the data provided insufficient signal to reliably estimate gene flow, as indicated by the posterior distributions consistently following the prior distributions (Table S3)..

### 3.3 High discordance in other members of South Asian clade

The gCF and sCF analysis across other members of SA clade, revealed no clear majority of gene trees (mostly showing high gene discordance factor due to polyphyly) or sites supporting a single topology, indicating extensive genealogical discordance (Fig. 4A-B; Fig. S14-S21).

### 3.4 Poor phylogenetic resolution of plastid markers

Two plastid markers (*trnL-LF* and partial *matK*) amplification and sequencing for Indian *Ledebouria* samples (Table 1) were done to check if they were also able to capture similar phylogenetic resolution as Angiosperms353 nuclear markers (see methods).

The plastid *trnL-LF* dataset had 70 taxa and 1145 aligned characters, of which 187 were parsimony-informative. Bayesian phylogenetic analysis did not retrieve the SA clade as monophyletic (Fig. 5). Nevertheless, the four clades observed in Howard et al. (2022) were loosely retrieved and their interrelationships remained unresolved (Fig. 5). Indian accessions from the present study were placed within *Ledebouria* clade B, as expected (Fig. 5). This clade was well supported but included both our samples and African and Malagasy representatives of *Ledebouria* (Fig. 5). Several previously published GenBank sequences retained the name *L. revoluta*, reflecting their publication prior to Chakral et al. (2024). The *trnL-LF* marker was able to distinguish true (African) *L. revoluta* from Indian entities—for example, the accession Nordal and Stedje 2409, which was recovered in the paraphyletic *Ledebouria* clade A (Fig. 5).

**Fig. 5.**
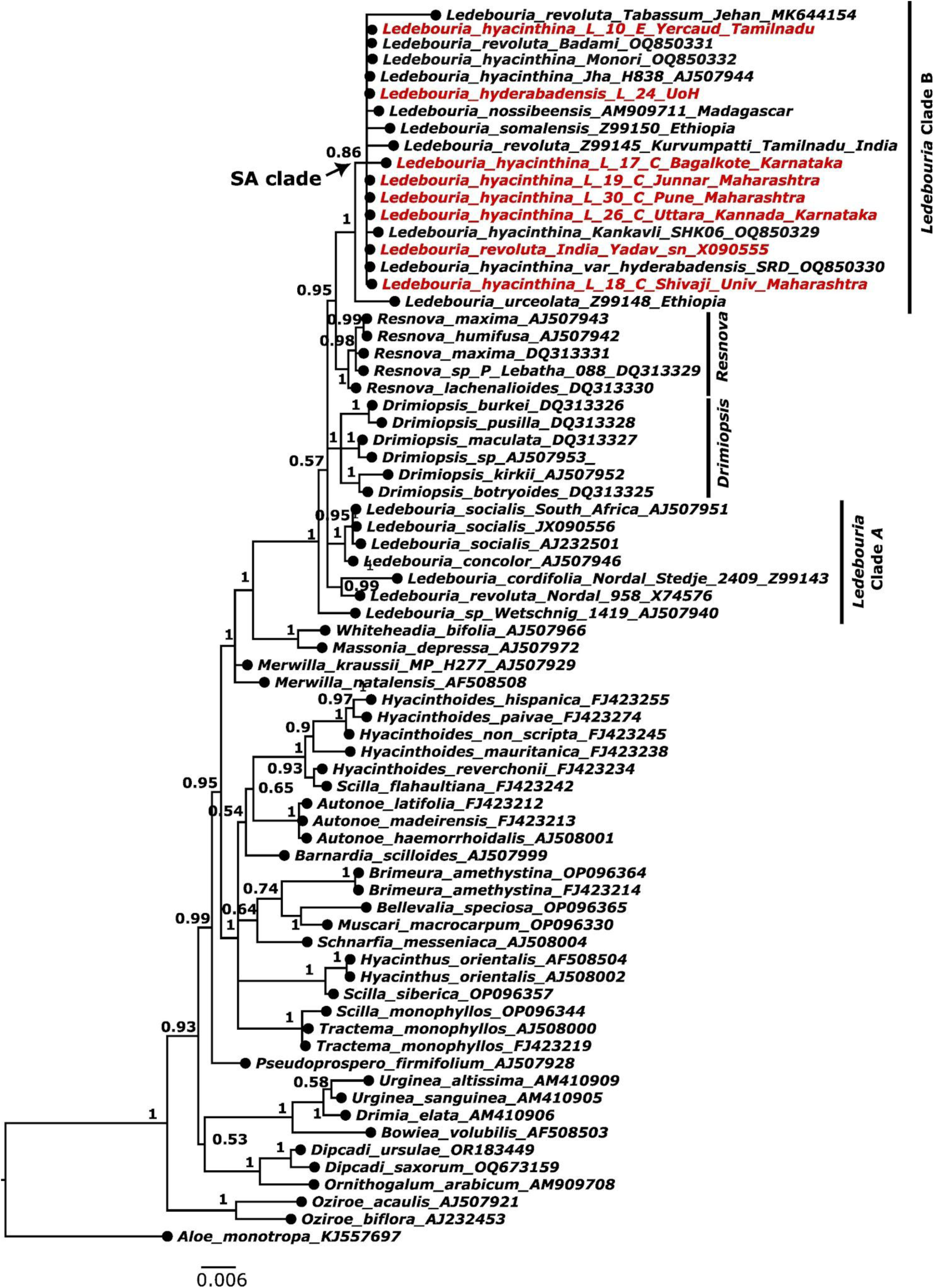
Bayesian phylogeny analysis of the *trnL*-*LF* dataset. Sequences generated for this study are indicated in red. Numbers on branches indicate posterior probability values.

The *trnL-LF* dataset lacked sufficient phylogenetic resolution to distinguish Indian *Ledebouria* from African taxa, with these accessions collapsing into a polytomy within *Ledebouria* clade B (Fig. 5). In this analysis, the *Resnova* clade was recovered as sister to *Ledebouria* clade B, in contrast to the Angiosperms353 dataset, where it was resolved as sister to *Ledebouria* clade A (Fig. 1-3; 5).

The plastid partial *matK* dataset included 19 sequences generated in this study and comprised 42 taxa with 905 aligned characters, of which only 91 were parsimony-informative. Bayesian phylogenetic analysis yielded poorly resolved relationships among clades (Fig. S24). Nonetheless, the *matK* dataset was able to distinguish the true (African) *L. revoluta* (accession Van Neste KM 688) from the Indian entities (Fig. S24), now recognized as *L. hyacinthina* (Chakral et al., 2024).

## 4. Discussion

The main goal of the study was to understand species boundaries within Indian *Ledebouria* using molecular phylogenetics with a robust set of DNA markers. The first molecular phylogenetic analysis of Indian *Ledebouria* was published by Chakral et al. (2024). Their work clarified the distinction between African *L*. *revoluta* and the Indian entities of the same name and hence, they resurrected the name *L*. *hyacinthina* (Roth) from the former synonym *S. hyacinthina* (Roth) Macbride (Chakral et al., 2024). They used two plastid DNA regions—the *trnC-ycf6* intron and the *trnL-trnF* intergenic spacer—to reconstruct the phylogeny. However, due to the unavailability of sequences, African *L*. *revoluta* could not be included in their *trnC*-*ycf6* analysis. In their *trnL*-*trnF* analysis, *Ledebouria* was found to be polyphyletic, with taxa appearing in two distantly related clades. Although current understanding supports the polyphyly of *Ledebouria*, with *Resnova* and *Drimiopsis* nested within it (Howard et al., 2022; Lebatha et al., 2006), the results of Chakral et al. (2024) differ—possibly due to an inappropriate choice of outgroup and the limited phylogenetic signal from using only the intergenic spacer. In contrast, our analysis, which includes both the *trnL* intron and the *trnL-trnF* spacer, recovers *Ledebouria* sensu lato as monophyletic, including *Drimiopsis* and *Resnova*. This monophyly is also supported by our *matK* analysis. Nevertheless, both Chakral et al. (2024) and our plastid-based analyses show limited resolution among the Indian taxa.

In India, *Ledebouria* is widespread in the peninsular region’s grasslands. There are several minor morphological variants as well as karyotypic variants and a clear species delimitation is lacking. Especially there are different 2n chromosome counts such as 45, 60 and 90 which are several ploidy levels of the basic number 30 (Deshmukh, 2023; Giranje et al., 2019). The commonly used nuclear marker, the nuclear ribosomal internal transcribed spacer (ITS), could not be amplified with the universal primers (unpublished data), prompting our use of the Angiosperms353 nuclear marker set. With the publication of Howard et al. (2022), we had the opportunity to apply this marker set to global *Ledebouria* accessions. Our results combined with the data of Howard et al. (2022) yielded a robust phylogeny in which taxa were placed in expected clades—Indian accessions grouped with other Indian and Sri Lankan samples from their study, and our *Drimiopsis* sample in the expected *Drimiopsis* clade of their study.

Based on the various analyses performed in this study such as species tree inference, concordance analysis and species delimitation, it seems *L. hyacinthina* var. *hyderabadensis* might be a putative species but a follow-up study with better sampling shoukd be done for confirmation. In addition it exhibits the most distinct morphology among Indian accessions, characterized by fewer than 25 flowers per inflorescence (compared to at least 40 in others), presence of sterile flowers, and a beaked ovary—traits highlighted by Chakral et al. (2024). These morphological distinctions are supported by our molecular analysis using the Angiosperms353 nuclear marker set, suggesting that *L. hyacinthina* var. *hyderabadensis* should be elevated to species rank.

In contrast, the remaining accessions of *L. hyacinthina* show no significant molecular sequence differentiation, despite minor morphological variation at the population level. A prior study on male meiosis across 18 populations revealed three cytotypes (n = 15, 30, 45), with mostly normal meiosis, high pollen viability, and uniform pollen morphology, indicating no correlation between chromosome number and reproductive fitness (Deshmukh et al., 2024). The phylogenomic discordance observed in the remaining accessions of *L. hyacinthina* might be due to limited phylogenetic information, gene-tree estimation error, incomplete lineage sorting and high gene flow. Further analyses with dense sampling is required for better resolution of their evolutionary history.

Species complexes arise when overlapping morphological characters form a continuum of phenotypic variation, preventing clear taxonomic delimitation (Grant, 1981). Such complexes resist resolution by conventional methods, necessitating integrative approaches like phylogenetic reconstruction, biogeographic analysis, or environmental niche modelling (Joshi and Karanth, 2012). Widespread species, especially within monocots, exhibit species complexes (Ashokan and Gowda, 2026; Surveswaran et al., 2018) and an integrative approach is often sought to unravel such species complexes. In our analysis, even a large molecular marker set failed to resolve *Ledebouria* into distinct geographical or karyotypic clades, revealing a widespread lineage in South Asia with extensive cryptic diversity and an unresolved phylogeny. This study represents only a preliminary insight into the diversity of this lineage in peninsular India. A limitation is our sparse taxon sampling, although it was designed to represent distinct karyotypes. Future work incorporating denser sampling and a more extensive marker set — such as the Asparagaceae1726 probe set (Bentz and Leebens-Mack, 2024) — may better resolve the evolutionary patterns within this group.

## Data availability statement

The sequences generated in this study have been deposited in NCBI with Bioproject ID: PRJNA1558098.

The supercontigs sequences of the samples used in this study are attached as supplementary material along with this publication (Data S1).

## CRediT authorship contribution statement

Sachidanand Nayak: Formal analysis, Writing - Original Draft, Data Curation, Investigation. Pradip Deshmukh: Resources, Data curation, Investigation. S.R. Yadav: Resources, Data curation, Investigation, Writing - Review & Editing. Manoj M. Lekhak: Resources, Data curation, Investigation, Writing - Review & Editing. Siddharthan Surveswaran: Conceptualization, Methodology, Project administration, Funding acquisition, Investigation, Writing - Original Draft, Data Curation.

## Author’s statement

All authors have read the final draft and agree to the publication, ensuring that all individuals deserving authorship have been appropriately acknowledged in the authorship contribution statement.

## Declaration of competing interest

The authors declare that they have no known competing financial interests or personal relationships that could have appeared to influence the work reported in this paper.

## Acknowledgements

Siddharthan Surveswaran thanks the University of Hyderabad Institution of Eminence grant number UoH-IoE-RC5-22-042 for funding.

## Supplementary data

Fig. S1. Species tree generated by ASTRAL of *Ledebouria* taxa using the untrimmed dataset containing 334 genes. Numbers at nodes indicate local posterior probability (LPP) support. Sequences generated for this study are indicated in red.

Fig. S2. Species tree generated by ASTRAL of *Ledebouria* taxa using the untrimmed dataset containing 106 genes. Numbers at nodes indicate local posterior probability (LPP) support. Sequences generated for this study are indicated in red.

Fig. S3. Species tree generated by ASTRAL of *Ledebouria* taxa using the trimmed dataset containing 106 genes. Numbers at nodes indicate local posterior probability (LPP) support. Sequences generated for this study are indicated in red.

Fig. S4. Species tree generated by IQTREE of *Ledebouria* taxa using the untrimmed dataset containing 334 genes. Numbers at nodes indicate SH-aLRT (SH approximate likelihood ratio tests) and ultrafast bootstrap values. Sequences generated for this study are indicated in red.

Fig. S5. Species tree generated by IQTREE of *Ledebouria* taxa using the untrimmed dataset containing 106 genes. Numbers at nodes indicate SH-aLRT (SH approximate likelihood ratio tests) and ultrafast bootstrap values. Sequences generated for this study are indicated in red.

Fig. S6. Species tree generated by IQTREE of *Ledebouria* taxa using the trimmed dataset containing 106 genes. Numbers at nodes indicate SH-aLRT (SH approximate likelihood ratio tests) and ultrafast bootstrap values. Sequences generated for this study are indicated in red.

Fig. S7. Weighted ASTRAL generated species tree of *Ledebouria* taxa using the untrimmed dataset containing 334 genes. Numbers at nodes indicate local posterior probability (LPP) support. Sequences generated for this study are indicated in red.

Fig. S8. Weighted ASTRAL generated species tree of *Ledebouria* taxa using the trimmed dataset containing 334 genes. Numbers at nodes indicate local posterior probability (LPP) support. Sequences generated for this study are indicated in red.

Fig. S9. Weighted ASTRAL generated species tree of *Ledebouria* taxa using the untrimmed dataset containing 106 genes. Numbers at nodes indicate local posterior probability (LPP) support. Sequences generated for this study are indicated in red.

Fig. S10. Asteroid generated species tree of *Ledebouria* taxa using the untrimmed dataset containing 334 genes. Numbers at nodes indicate bootstrap support. Sequences generated for this study are indicated in red.

Fig. S11. Asteroid generated species tree of *Ledebouria* taxa using the untrimmed dataset containing 106 genes. Numbers at nodes indicate bootstrap support. Sequences generated for this study are indicated in red.

Fig. S12. Asteroid generated species tree of *Ledebouria* taxa using the trimmed dataset containing 334 genes. Numbers at nodes indicate bootstrap support. Sequences generated for this study are indicated in red.

Fig. S13. Asteroid generated species tree of *Ledebouria* taxa using the trimmed dataset containing 106 genes. Numbers at nodes indicate bootstrap support. Sequences generated for this study are indicated in red.

Fig. S14. NNI-1 species tree for the branch leading to SA clade. Sequences generated for this study are indicated in red.

Fig. S15. NNI-2 species tree for the branch leading to SA clade. Sequences generated for this study are indicated in red.

Fig. S16. Gene concordance factor analysis of the *Ledebouria* SA clade using the untrimmed dataset containing 334 genes. Numbers on the nodes indicate Gene concordance factor/Gene discordance factor for Nearest Neighbour Interchange (NNI)-1 branch/Gene discordance factor for NNI-2 branch/Gene discordance factor due to polyphyly. Sequences generated for this study are indicated in red.

Fig. S17. Gene concordance factor analysis of the *Ledebouria* SA clade using the trimmed dataset containing 334 genes. Numbers on the nodes indicate Gene concordance factor/Gene discordance factor for Nearest Neighbour Interchange (NNI)-1 branch/Gene discordance factor for NNI-2 branch/Gene discordance factor due to polyphyly. Sequences generated for this study are indicated in red.

Fig. S18. Gene concordance factor analysis of the *Ledebouria* SA clade using the trimmed dataset containing 106 genes. Numbers on the nodes indicate Gene concordance factor/Gene discordance factor for Nearest Neighbour Interchange (NNI)-1 branch/Gene discordance factor for NNI-2 branch/Gene discordance factor due to polyphyly. Sequences generated for this study are indicated in red.

Fig. S19. Site concordance factor analysis of the *Ledebouria* SA clade using the untrimmed dataset containing 334 genes. Numbers on the nodes indicate Site concordance factor averaged over 100 quartets/Site discordance factor for alternative quartet 1/Site discordance factor for alternative quartet 2. Sequences generated for this study are indicated in red.

Fig. S20. Site concordance factor analysis of the *Ledebouria* SA clade using the trimmed dataset containing 334 genes. Numbers on the nodes indicate Site concordance factor averaged over 100 quartets/Site discordance factor for alternative quartet 1/Site discordance factor for alternative quartet 2. Sequences generated for this study are indicated in red.

Fig. S21. Site concordance factor analysis of the *Ledebouria* SA clade using the trimmed dataset containing 106 genes. Numbers on the nodes indicate Site concordance factor averaged over 100 quartets/Site discordance factor for alternative quartet 1/Site discordance factor for alternative quartet 2. Sequences generated for this study are indicated in red.

Fig. S22. Barplots showing the frequency of each model evaluated during species delimitation analysis. (A) and (B): thetaprior=invgamma 3 0.002 int and tauprior=invgamma 3 0.01; (C) and (D): thetaprior=invgamma 3 0.005 int and tauprior=invgamma 3 0.03; (E) and (F): thetaprior=invgamma 3 0.01 int and tauprior=invgamma 3 0.06.

Fig. S23. Lineplots showing the distribution of *lnL* across iteration for each model evaluated during species delimitation analysis. (A) and (B): thetaprior=invgamma 3 0.002 int and tauprior=invgamma 3 0.01; (C) and (D): thetaprior=invgamma 3 0.005 int and tauprior=invgamma 3 0.03; (E) and (F): thetaprior=invgamma 3 0.01 int and tauprior=invgamma 3 0.06.

Fig. S24. Bayesian phylogeny analysis of *matK* data. Sequences generated for this study are indicated in red.

Table S1: Hybpiper assembly statistics

Table S2: Results of species delimitation analysis. Each digit of the model represents an ancestral node. 0 and 1 in the model represents collapse and retention of the corresponding ancestral node respectively. Three different set of theta and tau prior were used. Each prior set analysis were repeated twice with random system generated seed values. The models explored in each replicate of each prior set are highlighted in yellow.

Table S3: Results of gene flow analysis. Node 1=*Ledebouria revoluta*, Node 2= *Ledbouria hyacinthina* and Node 3=*Ledbouria hyderabenesis*

Data S1: Assembled supercontigs of all samples used in this study, generated using Angiosperms353 target capture sequencing, together with control and input files used for species delimitation and gene flow analyses.

## Notes

### Competing Interest Statement

The authors have declared no competing interest.

### Summary of Updates

The species delimitation and gene flow analyses had been updated. Additional phylogenetic analysis was performed.

## References

Ashokan, A., Gowda, V., 2026. Things aren’t always black or white — Phylogenetic systematics of ginger-lilies with insights into species complexes. Taxon 75. 10.1002/tax.70081.

Baker, J.G., 1896. *Scilla*, in: Thiselton-Dyer, W.T. (Ed.), Flora Capensis :Being a Systematic Description of the Plants of the Cape Colony, Caffraria, & Port Natal (and Neighbouring Territories). L. Reeve and Company, London, pp. 478–494.

Baker, J.G., 1872. Revision of the genera and species of Scilleae and Chlorogaleae. Journal of the Linnean Society of London, Botany 13, 209–266. 10.1111/j.1095-8339.1872.tb00093.x.

Bentz, P.C., Leebens-Mack, J., 2024. Developing Asparagaceae1726: An Asparagaceae-specific probe set targeting 1726 loci for Hyb-Seq and phylogenomics in the family. Appl. Plant Sci. 10.1002/aps3.11597.

Buerki, S., Jose, S., Yadav, S.R., Goldblatt, P., Manning, J.C., Forest, F., 2012. Contrasting biogeographic and diversification patterns in two Mediterranean-type ecosystems. PLoS ONE 7, e39377. 10.1371/journal.pone.0039377.

Capella-Gutiérrez, S., Silla-Martínez, J.M., Gabaldón, T., 2009. trimAl: A tool for automated alignment trimming in large-scale phylogenetic analyses. Bioinformatics 25, 1972–1973. 10.1093/bioinformatics/btp348.

Chakral, K.G., Dutta, S.R., Rodrigues, H., Dwivedi, M., 2024. Revisionary insights into the genus *Ledebouria* (Asparagaceae) from India. Phytotaxa 641, 1–21. 10.11646/phytotaxa.641.1.1.

Cuénoud, P., Savolainen, V., Chatrou, L.W., Powell, M., Grayer, R.J., Chase, M.W., 2002. Molecular phylogenetics of Caryophyllales based on nuclear 18S rDNA and plastid rbcL, atpB, and matK DNA sequences. Am. J. Bot. 89, 132–144. 10.3732/ajb.89.1.132.

Darriba, D., Taboada, G.L., Doallo, R., Posada, D., 2012. jModelTest 2: more models, new heuristics and parallel computing. Nat. Methods 9, 772. 10.1038/nmeth.2109.

Deshmukh, P.V., Yadav, S.R., Lekhak, M.M., 2024. Male meiosis and pollen morphology of some populations of *Ledebouria revoluta* (Asparagaceae) from India. Cytologia (Tokyo) 89, 97–104. 10.1508/cytologia.89.97.

Deshmukh, P.V., 2023. Biosystematic Studies In *Ledebouria revoluta* (Asparagaceae) In India (Doctoral dissertation). Shivaji University Kolhapur.

Dixit, G.B., Yadav, S.R., Salunkhe, C.B., 1989. Cytomorphological studies in Scilla hyacinthina (Roth) Macbr. complex from Maharashtra., in: Proc Conf on Cytol Genet. pp. 124–134.

Flouri, T., Jiao, X., Huang, J., Rannala, B., Yang, Z., 2023. Efficient Bayesian inference under the multispecies coalescent with migration. Proc Natl Acad Sci USA 120, e2310708120. 10.1073/pnas.2310708120.

Flouri, T., Jiao, X., Rannala, B., Yang, Z., 2018. Species tree inference with BPP using genomic sequences and the multispecies coalescent. Mol. Biol. Evol. 35, 2585–2593. 10.1093/molbev/msy147.

Ford, C.S., Ayres, K.L., Toomey, N., Haider, N., Van Alphen Stahl, J., Kelly, L.J., Wikström, N., Hollingsworth, P.M., Duff, R.J., Hoot, S.B., Cowan, R.S., Chase, M.W., Wilkinson, M.J., 2009. Selection of candidate coding DNA barcoding regions for use on land plants. Botan. J. Linn. Soc. 159, 1–11. 10.1111/j.1095-8339.2008.00938.x.

Gadagkar, S.R., Rosenberg, M.S., Kumar, S., 2005. Inferring species phylogenies from multiple genes: concatenated sequence tree versus consensus gene tree. J. Exp. Zool. B Mol. Dev. Evol. 304, 64–74. 10.1002/jez.b.21026.

Giranje, P.T., Mayur D., N.D., Yadav, S.R., 2019. A Karyotype analysis in *Ledebouria revoluta* (Hyacinthaceae) with a new Cytotype. Chromosome Botany 13, 61–67. 10.3199/iscb.13.61.

Giranje, P.T., Nandikar, M.D., 2016. Synopsis of the genus *Ledebouria* Roth (Hyacinthaceae: Hyacinthoideae) in India. Webbia 71, 213–217. 10.1080/00837792.2016.1182324.

Grant, V., 1981. Plant Speciation. Columbia University Press. 10.7312/gran92318.

Guindon, S., Dufayard, J.-F., Lefort, V., Anisimova, M., Hordijk, W., Gascuel, O., 2010. New Algorithms and Methods to Estimate Maximum-Likelihood Phylogenies: Assessing the Performance of PhyML 3.0. Syst. Biol. 59, 307–321. 10.1093/sysbio/syq010.

Hoang, D.T., Chernomor, O., von Haeseler, A., Minh, B.Q., Vinh, L.S., 2018. Ufboot2: improving the ultrafast bootstrap approximation. Mol. Biol. Evol. 35, 518–522. 10.1093/molbev/msx281.

Howard, C.C., Crowl, A.A., Harvey, T.S., Cellinese, N., 2022. Peeling back the layers: First phylogenomic insights into the Ledebouriinae (Scilloideae, Asparagaceae). Mol. Phylogenet. Evol. 169, 107430. 10.1016/j.ympev.2022.107430.

Jessop, J.P., 1970. Studies in the Bulbous Liliaceae in South Africa: 1. Scilla, Schizocarpus and Ledebouria. Journal of South African Botany 36, 233–266.

Johnson, M.G., Gardner, E.M., Liu, Y., Medina, R., Goffinet, B., Shaw, A.J., Zerega, N.J.C., Wickett, N.J., 2016. HybPiper: Extracting coding sequence and introns for phylogenetics from high-throughput sequencing reads using target enrichment. Appl. Plant Sci. 4. 10.3732/apps.1600016.

Johnson, M.G., Pokorny, L., Dodsworth, S., Botigué, L.R., Cowan, R.S., Devault, A., Eiserhardt, W.L., Epitawalage, N., Forest, F., Kim, J.T., Leebens-Mack, J.H., Leitch, I.J., Maurin, O., Soltis, D.E., Soltis, P.S., Wong, G.K.-S., Baker, W.J., Wickett, N.J., 2019. A universal probe set for targeted sequencing of 353 nuclear genes from any flowering plant designed using k-medoids clustering. Syst. Biol. 68, 594–606. 10.1093/sysbio/syy086.

Joshi, J., Karanth, K.P., 2012. Coalescent method in conjunction with niche modeling reveals cryptic diversity among centipedes in the Western Ghats of South India. PLoS ONE 7, e42225. 10.1371/journal.pone.0042225.

Kalyaanamoorthy, S., Minh, B.Q., Wong, T.K.F., von Haeseler, A., Jermiin, L.S., 2017. ModelFinder: fast model selection for accurate phylogenetic estimates. Nat. Methods 14, 587–589. 10.1038/nmeth.4285.

Katoh, K., Rozewicki, J., Yamada, K.D., 2019. MAFFT online service: multiple sequence alignment, interactive sequence choice and visualization. Brief. Bioinformatics 20, bbx108. 10.1093/bib/bbx108.

Lanfear, R., Calcott, B., Ho, S.Y.W., Guindon, S., 2012. Partitionfinder: Combined selection of partitioning schemes and substitution models for phylogenetic analyses. Mol. Biol. Evol. 29, 1695–1701. 10.1093/molbev/mss020.

Lanfear, R., Calcott, B., Kainer, D., Mayer, C., Stamatakis, A., 2014. Selecting optimal partitioning schemes for phylogenomic datasets. BMC Evol. Biol. 14, 82. 10.1186/1471-2148-14-82.

Larsson, A., 2014. AliView: a fast and lightweight alignment viewer and editor for large datasets. Bioinformatics 30, 3276–3278. 10.1093/bioinformatics/btu531.

Lebatha, P., Buys, M.H., Stedje, B., 2006. *Ledebouria*, *Resnova* and *Drimiopsis*: A tale of three genera. Taxon 55, 643. 10.2307/25065640.

Li, H., 2013. Aligning sequence reads, clone sequences and assembly contigs with BWA-MEM. arXiv. 10.48550/arxiv.1303.3997.

Manning, J.C., Goldblatt, P., Fay, M.F., 2003. A revised generic synopsis of Hyacinthaceae in sub-Saharan Africa, based on molecular evidence, including new combinations and the new tribe Pseudoprospereae. Edin. Jnl of Bot. 60, 533–568. 10.1017/S0960428603000404.

Manning, J.C., Goldblatt, P., 2012. Hyacinthaceae. Bothalia 42, 47–48. 10.4102/abc.v42i1.27.

Manning, J.C., 2020. Systematics of *Ledebouria* sect. *Resnova* (Hyacinthaceae: Scilloideae: Massonieae), with a new subtribal classification of Massonieae. South African Journal of Botany 133, 98–110. 10.1016/j.sajb.2020.07.010.

Minh, B.Q., Hahn, M.W., Lanfear, R., 2020. New methods to calculate concordance factors for phylogenomic datasets. Mol. Biol. Evol. 37, 2727–2733. 10.1093/molbev/msaa106.

Morel, B., Williams, T.A., Stamatakis, A., 2023. Asteroid: a new algorithm to infer species trees from gene trees under high proportions of missing data. Bioinformatics 39. 10.1093/bioinformatics/btac832.

Pfosser, M., Wetschnig, W., Ungar, S., Prenner, G., 2003. Phylogenetic relationships among genera of Massonieae (Hyacinthaceae) inferred from plastid DNA and seed morphology. J. Plant Res. 116, 115–132. 10.1007/s10265-003-0076-8.

POWO, 2026. Ledebouria Roth | Plants of the World Online | Kew Science powo.science.kew.org [WWW Document]. URL https://powo.science.kew.org/taxon/urn:lsid:ipni.org:names:24421-1 (accessed 1.29.26).

Rambaut, A., Drummond, A.J., Xie, D., Baele, G., Suchard, M.A., 2018. Posterior summarization in Bayesian phylogenetics using Tracer 1.7. Syst. Biol. 67, 901–904. 10.1093/sysbio/syy032.

Rambaut, A., 2018. FigTree [Software], Version 1.4.4. Institute of Evolutionary Biology, University of Edinburgh.

Ronquist, F., Teslenko, M., van der Mark, P., Ayres, D.L., Darling, A., Höhna, S., Larget, B., Liu, L., Suchard, M.A., Huelsenbeck, J.P., 2012. MrBayes 3.2: efficient Bayesian phylogenetic inference and model choice across a large model space. Syst. Biol. 61, 539–542. 10.1093/sysbio/sys029.

Roth, A.W., 1821. Novae Plantarum Species praesertim Indiae Orientalis.

Seo, T.-K., Kishino, H., Thorne, J.L., 2005. Incorporating gene-specific variation when inferring and evaluating optimal evolutionary tree topologies from multilocus sequence data. Proc Natl Acad Sci USA 102, 4436–4441. 10.1073/pnas.0408313102.

Staden, R., Beal, K.F., Bonfield, J.K., 2000. The Staden package, 1998. Methods Mol. Biol. 132, 115–130. 10.1385/1-59259-192-2:115.

Stedje, B., 1998. Phylogenetic relationships and generic delimitation of sub-Saharan *Scilla* (Hyacinthaceae) and allied African genera as inferred from morphological and DNA sequence data. Plant Syst. Evol. 211, 1–11. 10.1007/BF00984908.

Surveswaran, S., Agarwala, K.P., Somani, S., Bhagyasree Raveendran, V.R., Vasudha, S., Toms, A., 2026. Out of South America into India: unusual long distance dispersal of a plant genus— *Lepidagathis* (Acanthaceae). J. Biogeogr. 53. 10.1111/jbi.70170.

Surveswaran, S., Gowda, V., Sun, M., 2018. Using an integrated approach to identify cryptic species, divergence patterns and hybrid species in Asian ladies’ tresses orchids (Spiranthes, Orchidaceae). Mol. Phylogenet. Evol. 124, 106–121. 10.1016/j.ympev.2018.02.025.

Tabatabaee, Y., Zhang, C., Warnow, T., Mirarab, S., 2023. Phylogenomic branch length estimation using quartets. Bioinformatics 39, i185–i193. 10.1093/bioinformatics/btad221.

Taberlet, P., Gielly, L., Pautou, G., Bouvet, J., 1991. Universal primers for amplification of three non-coding regions of chloroplast DNA. Plant Mol. Biol. 17, 1105–1109. 10.1007/BF00037152.

The Angiosperm Phylogeny Group, 2009. An update of the Angiosperm Phylogeny Group classification for the orders and families of flowering plants: APG III. Bot. J. Linn. Soc. 161, 105–121. 10.1111/j.1095-8339.2009.00996.x.

Wong, T.K.F., Ly-Trong, N., Ren, H., Baños, H., Roger, A., Susko, E., Bielow, C., De Maio, N., Goldman, N., Hahn, M., Huttley, G., Lanfear, R., Minh, B.Q., 2025. IQ-TREE 3: Phylogenomic Inference Software using Complex Evolutionary Models. 10.32942/X2P62N.

Zhang, C., Mirarab, S., 2022. Weighting by Gene Tree Uncertainty Improves Accuracy of Quartet-based Species Trees. Mol. Biol. Evol. 39. 10.1093/molbev/msac215.

Zhang, C., Nielsen, R., Mirarab, S., 2025. ASTER: A Package for Large-Scale Phylogenomic Reconstructions. Mol. Biol. Evol. 42. 10.1093/molbev/msaf172.

