## Supplementary Figures for "Species Delimitation in Indian *Ledebouria*: Resolving the putative species status of infra-generic taxon Despite Phylogenomic Discordance"

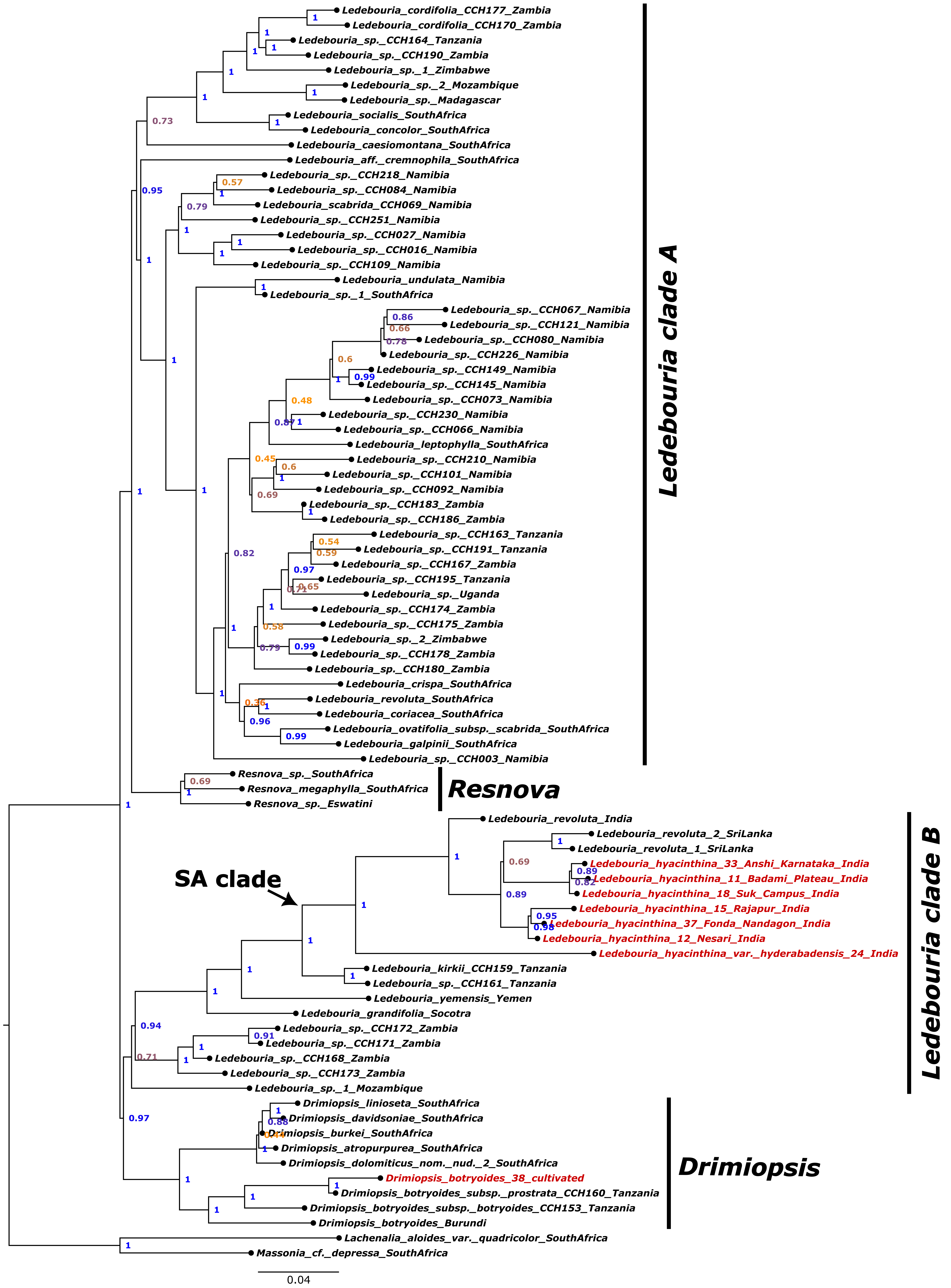


**Fig. S1. Species tree generated by ASTRAL of *Ledebouria* taxa using the untrimmed dataset containing 334 genes. Numbers at nodes indicate local posterior probability (LPP) support. Sequences generated for this study are indicated in red.**

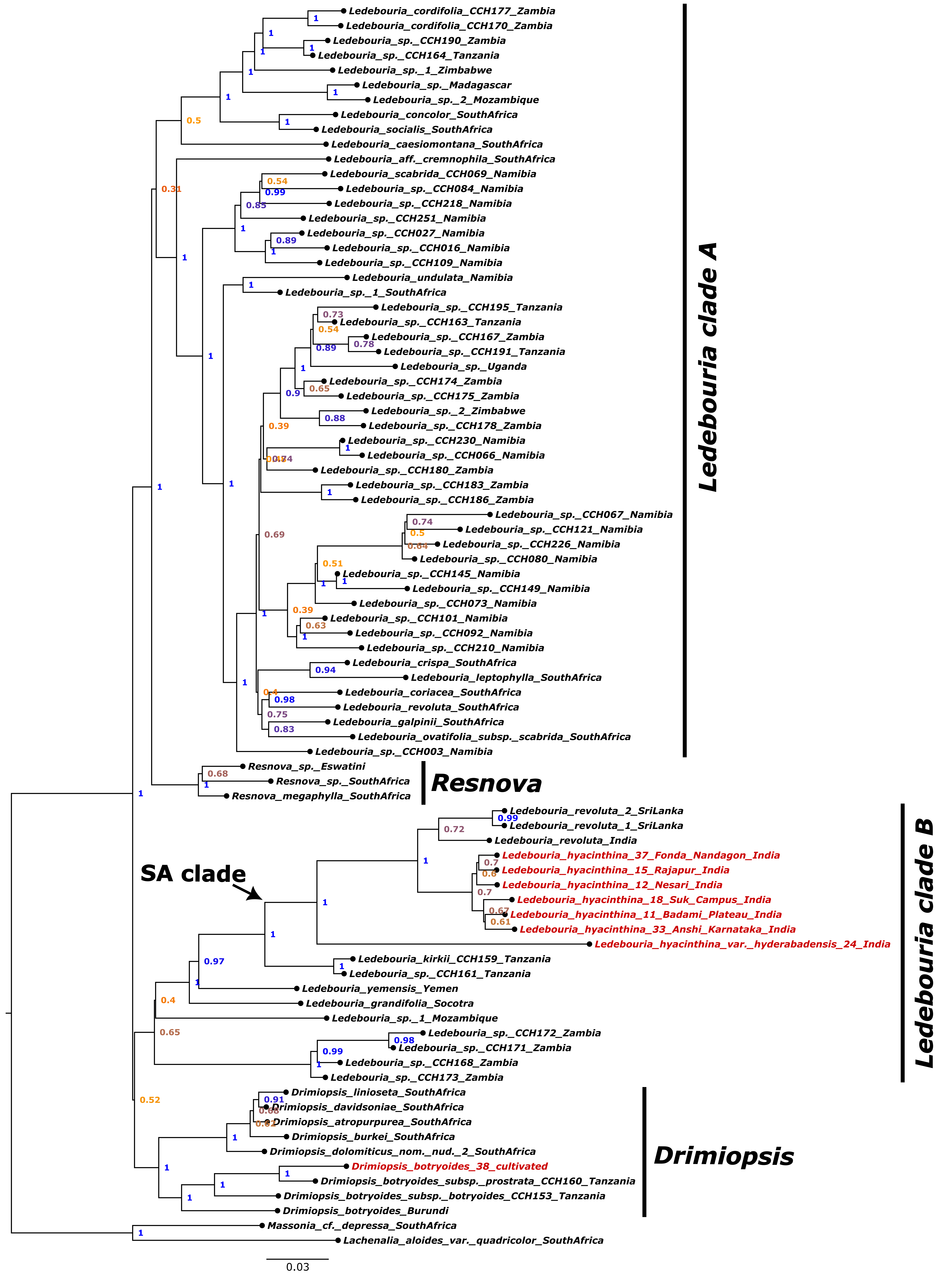


**Fig. S2. Species tree generated by ASTRAL of *Ledebouria* taxa using the untrimmed dataset containing 106 genes. Numbers at nodes indicate local posterior probability (LPP) support. Sequences generated for this study are indicated in red.**

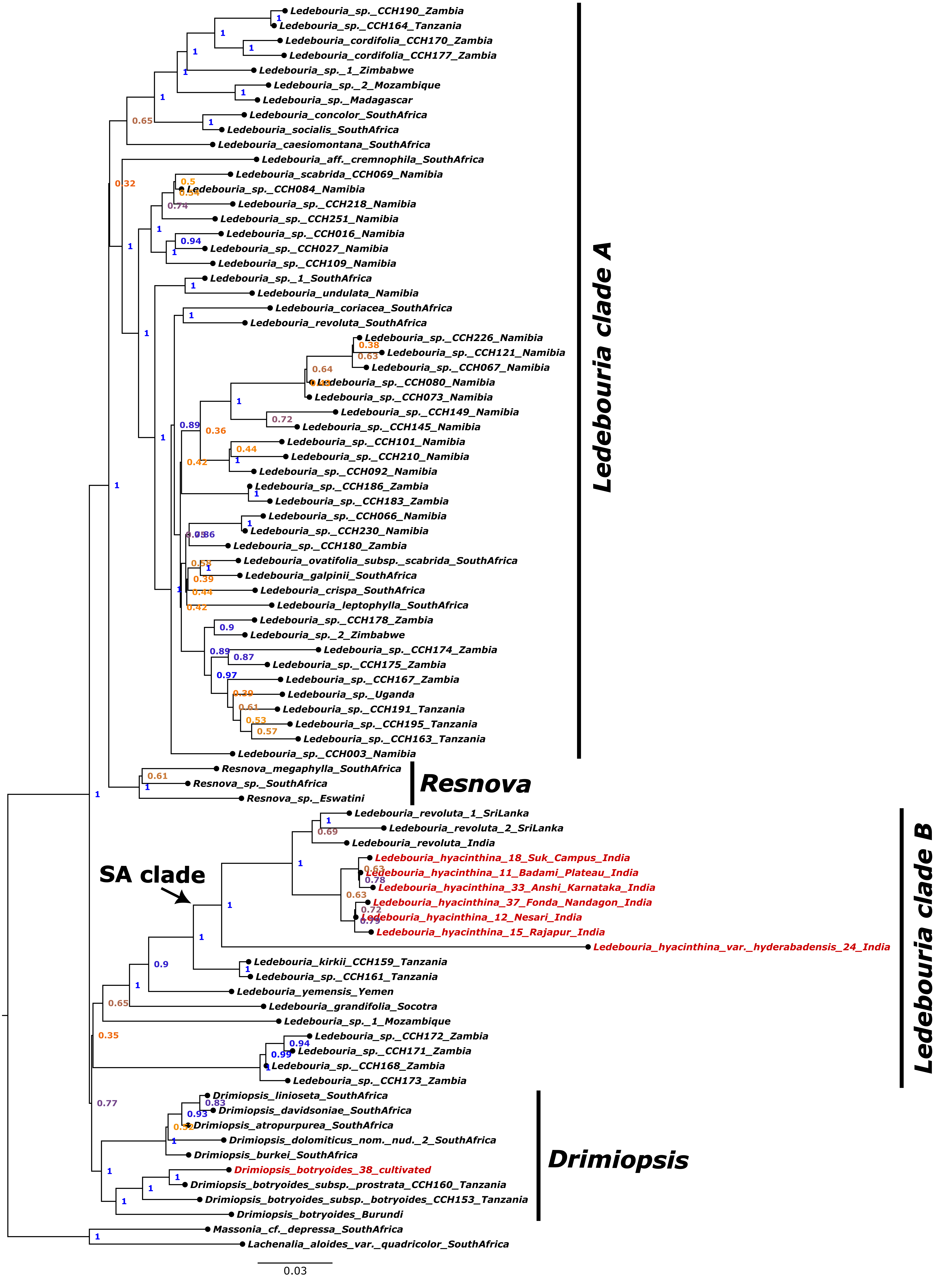


**Fig. S3. Species tree generated by ASTRAL of *Ledebouria* taxa using the trimmed dataset containing 106 genes. Numbers at nodes indicate local posterior probability (LPP) support. Sequences generated for this study are indicated in red.**

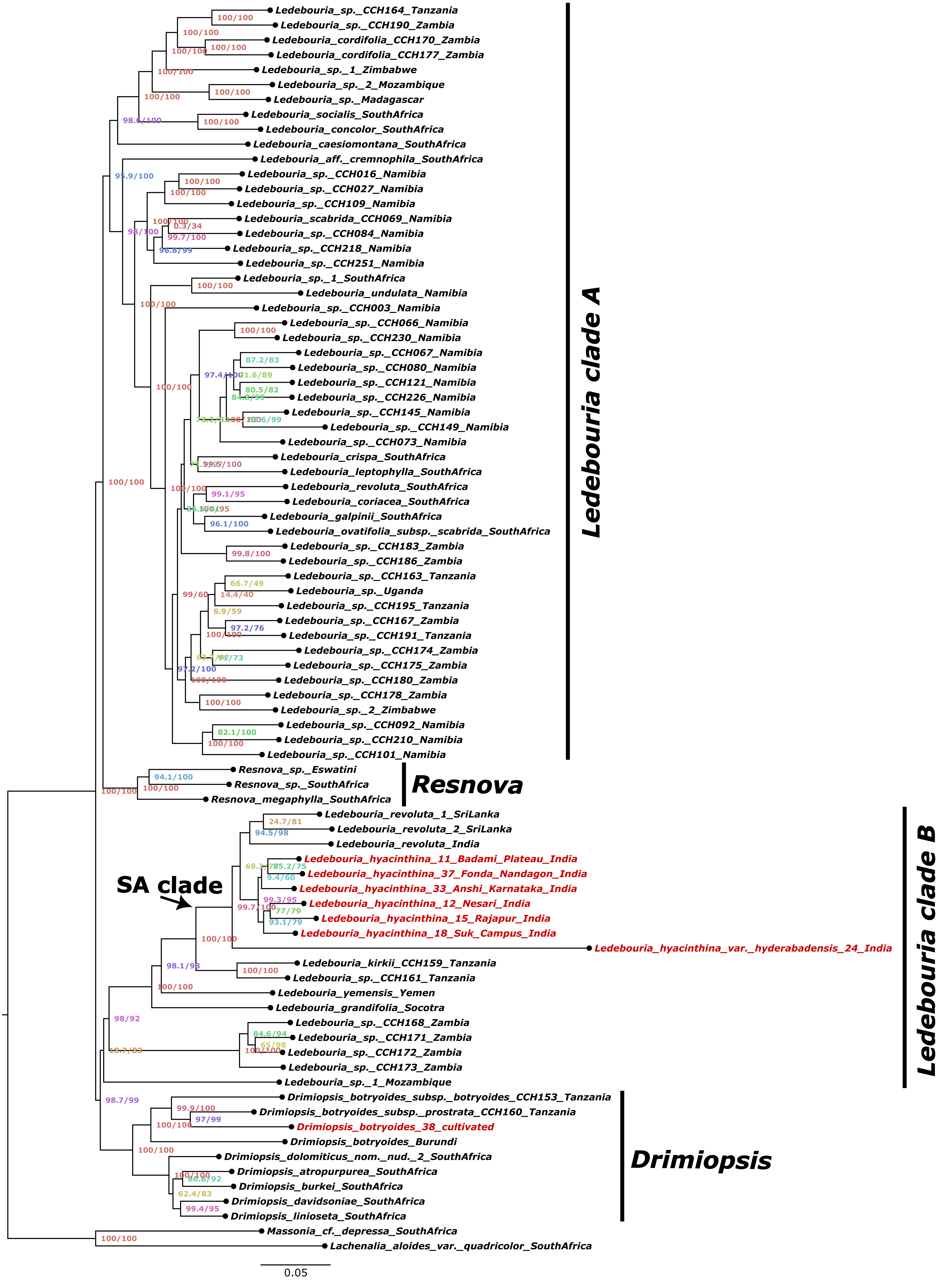


**Fig. S4. Species tree generated by IQTREE of *Ledebouria* taxa using the untrimmed dataset containing 334 genes. Numbers at nodes indicate SH-aLRT (SH approximate likelihood ratio tests) and ultrafast bootstrap values. Sequences generated for this study are indicated in red.**

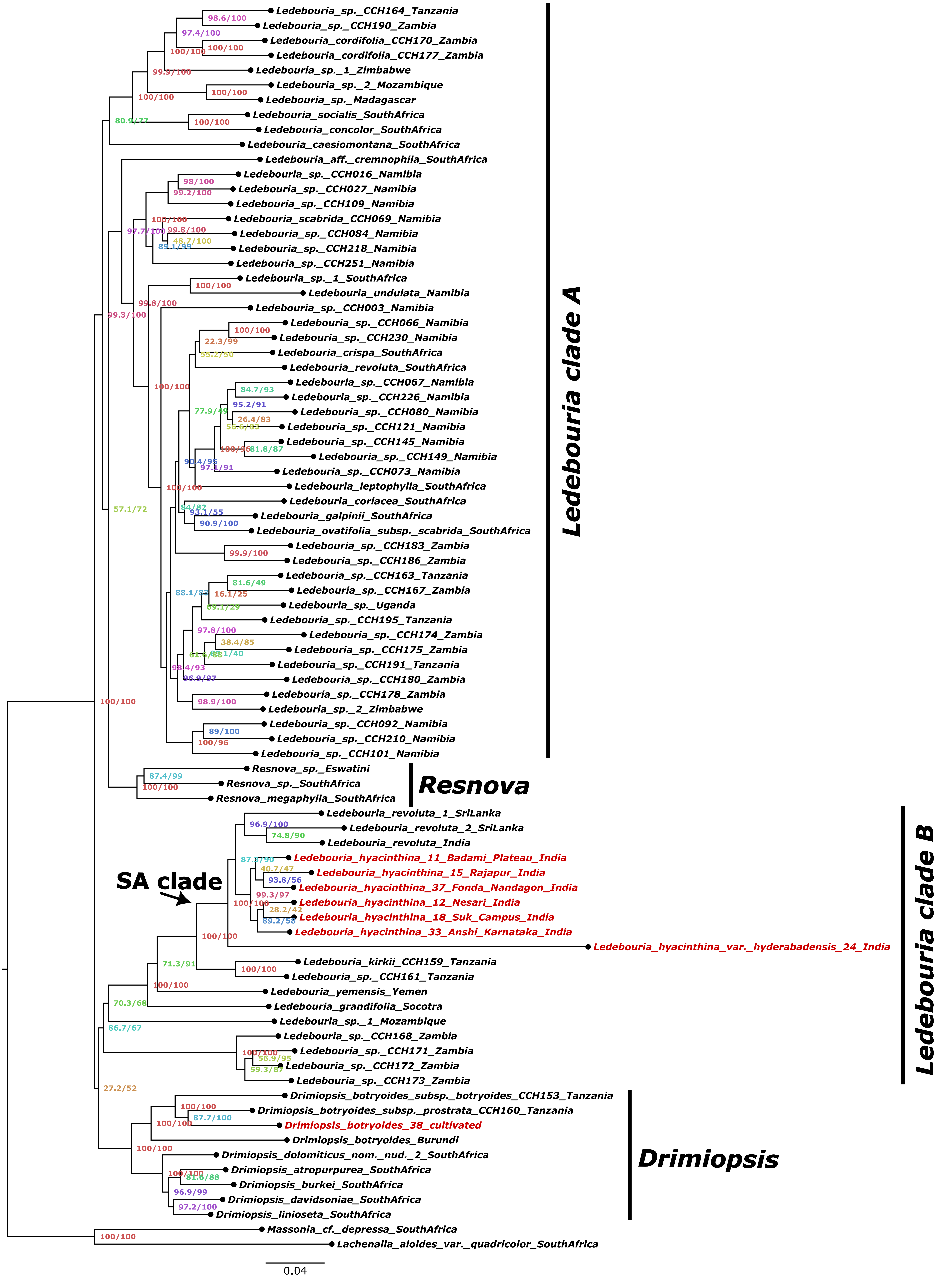


**Fig. S5. Species tree generated by IQTREE of *Ledebouria* taxa using the untrimmed dataset containing 106 genes. Numbers at nodes indicate SH-aLRT (SH approximate likelihood ratio tests) and ultrafast bootstrap values. Sequences generated for this study are indicated in red.**

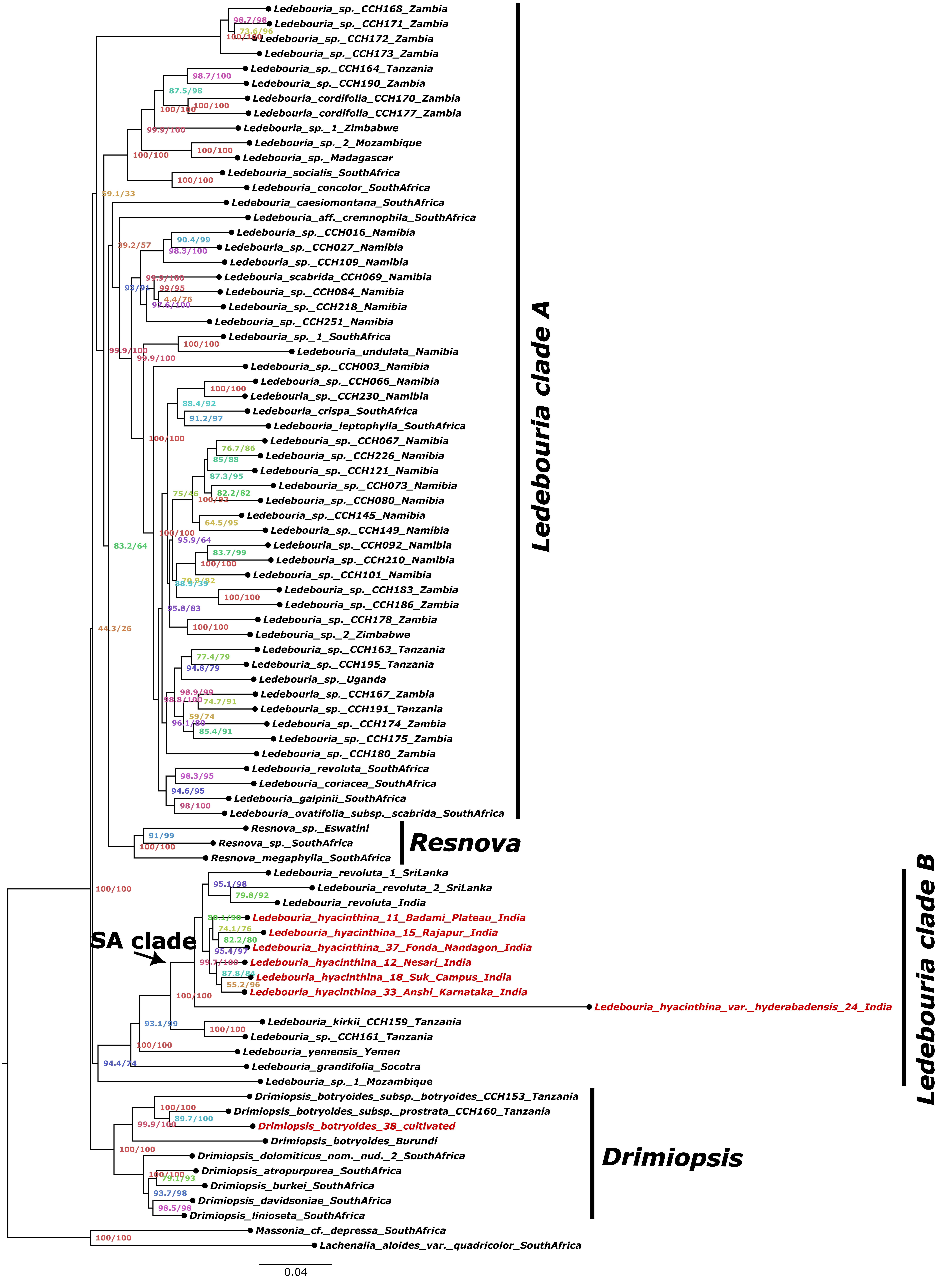


**Fig. S6. Species tree generated by IQTREE of *Ledebouria* taxa using the trimmed dataset containing 106 genes. Numbers at nodes indicate SH-aLRT (SH approximate likelihood ratio tests) and ultrafast bootstrap values. Sequences generated for this study are indicated in red.**

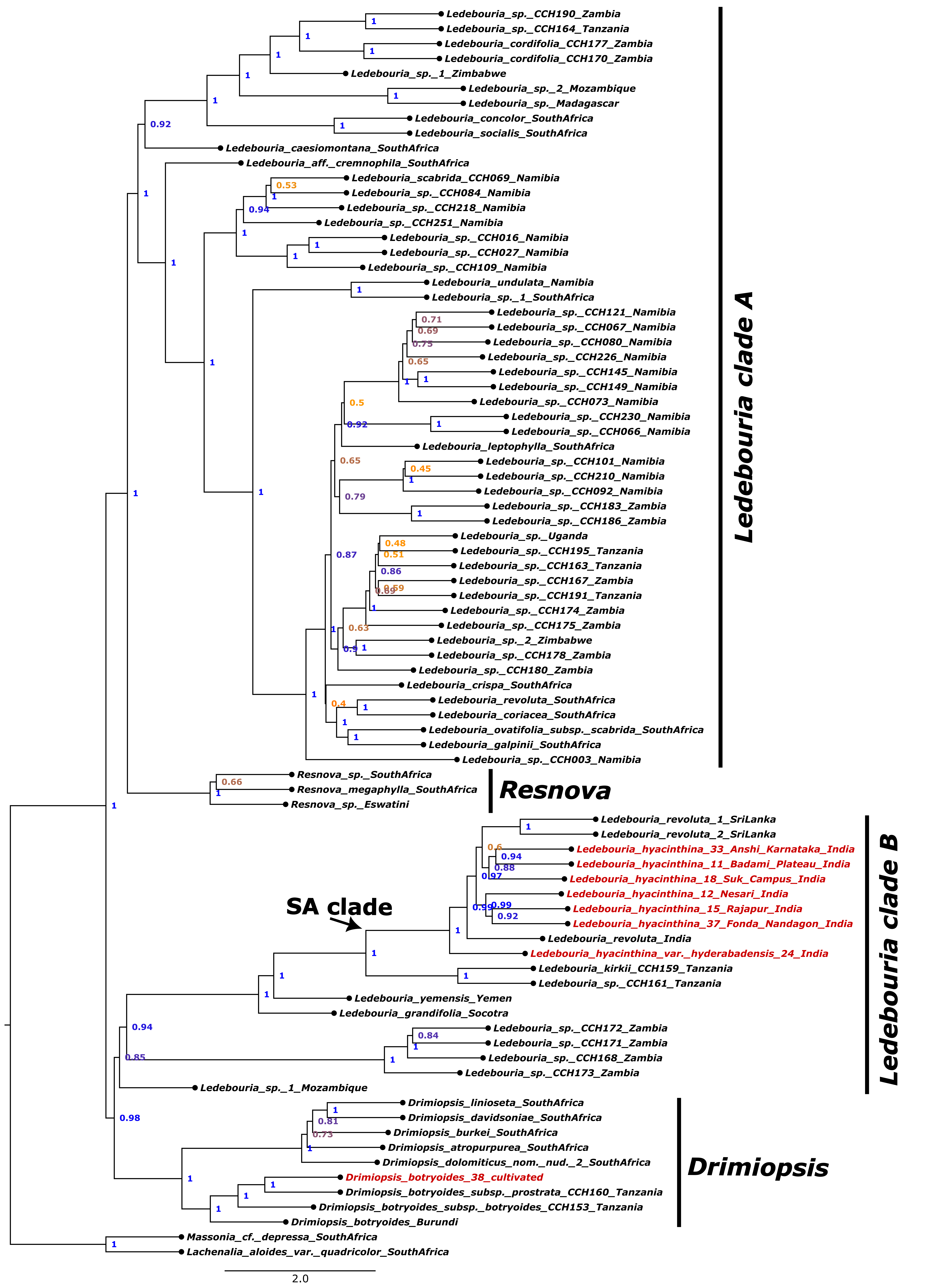


**Fig. S7. Weighted ASTRAL generated species tree of *Ledebouria* taxa using the untrimmed dataset containing 334 genes. Numbers at nodes indicate local posterior probability (LPP) support. Sequences generated for this study are indicated in red.**

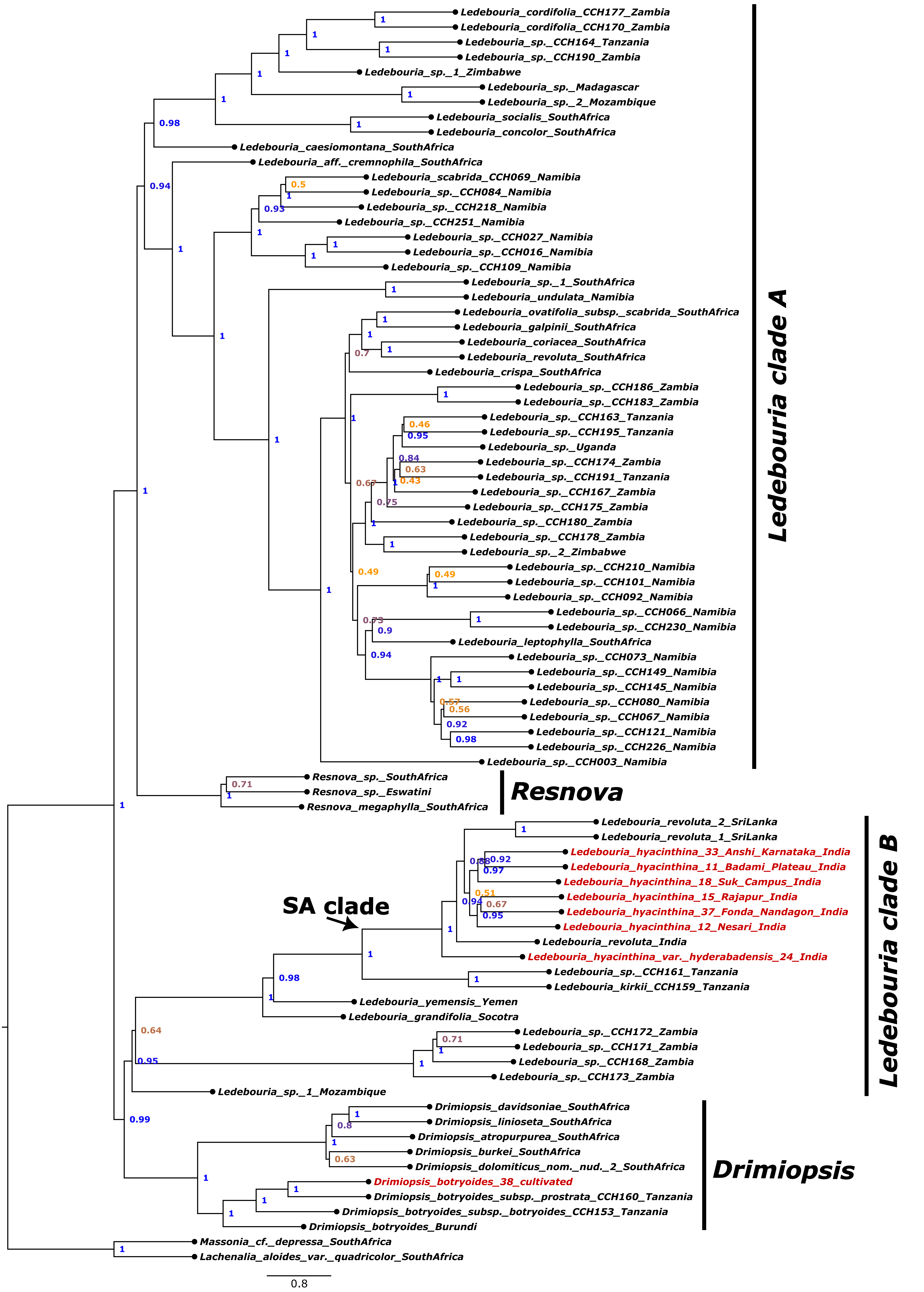


**Fig. S8. Weighted ASTRAL generated species tree of *Ledebouria* taxa using the trimmed dataset containing 334 genes. Numbers at nodes indicate local posterior probability (LPP) support. Sequences generated for this study are indicated in red.**

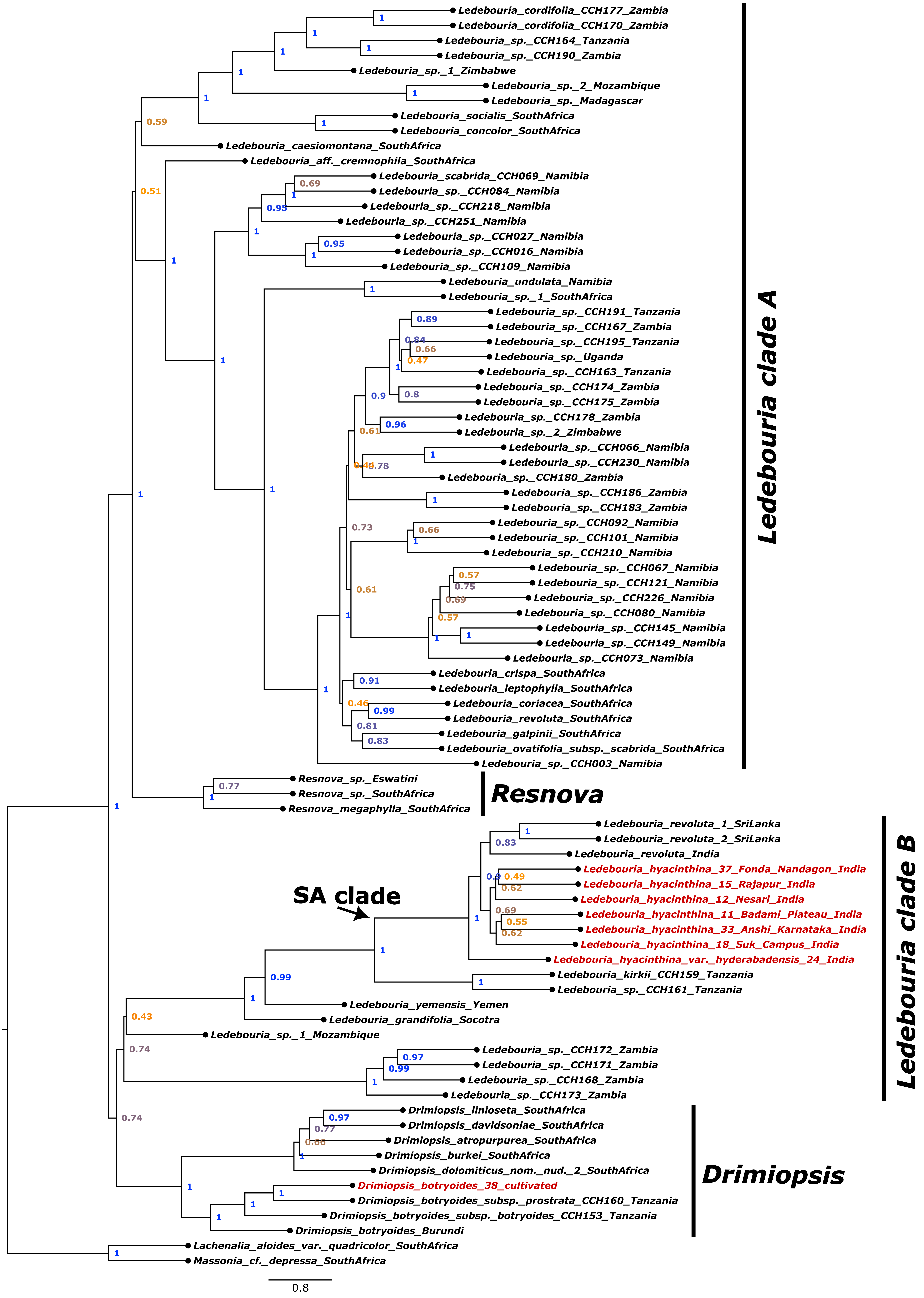


**Fig. S9.** Weighted ASTRAL generated species tree of *Ledebouria* taxa using the untrimmed dataset containing 106 genes. Numbers at nodes indicate local posterior probability (LPP) support. Sequences generated for this study are indicated in red.

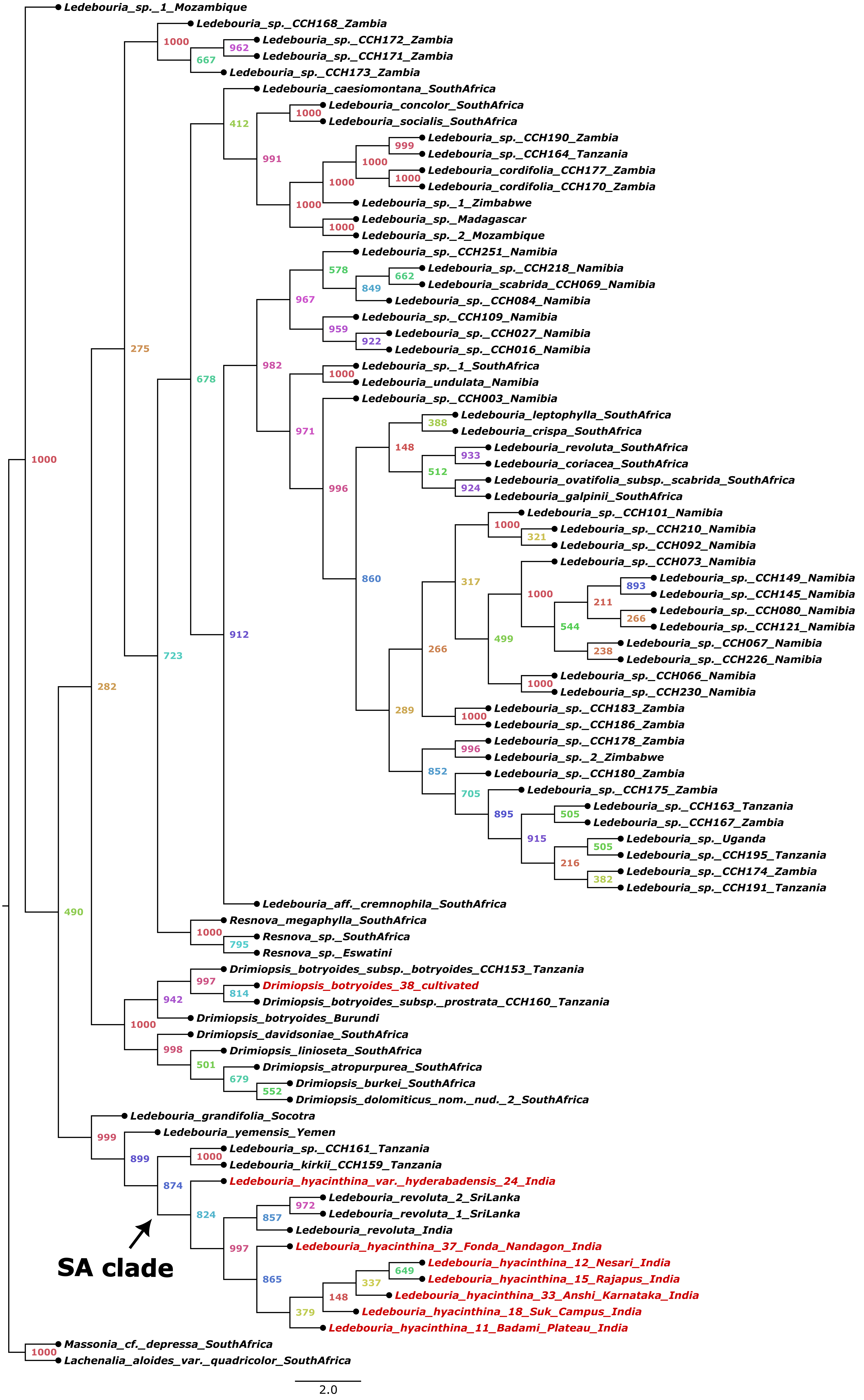


**Fig. S10. Asteroid generated species tree of *Ledebouria* taxa using the untrimmed dataset containing 334 genes. Numbers at nodes indicate bootstrap support. Sequences generated for this study are indicated in red.**

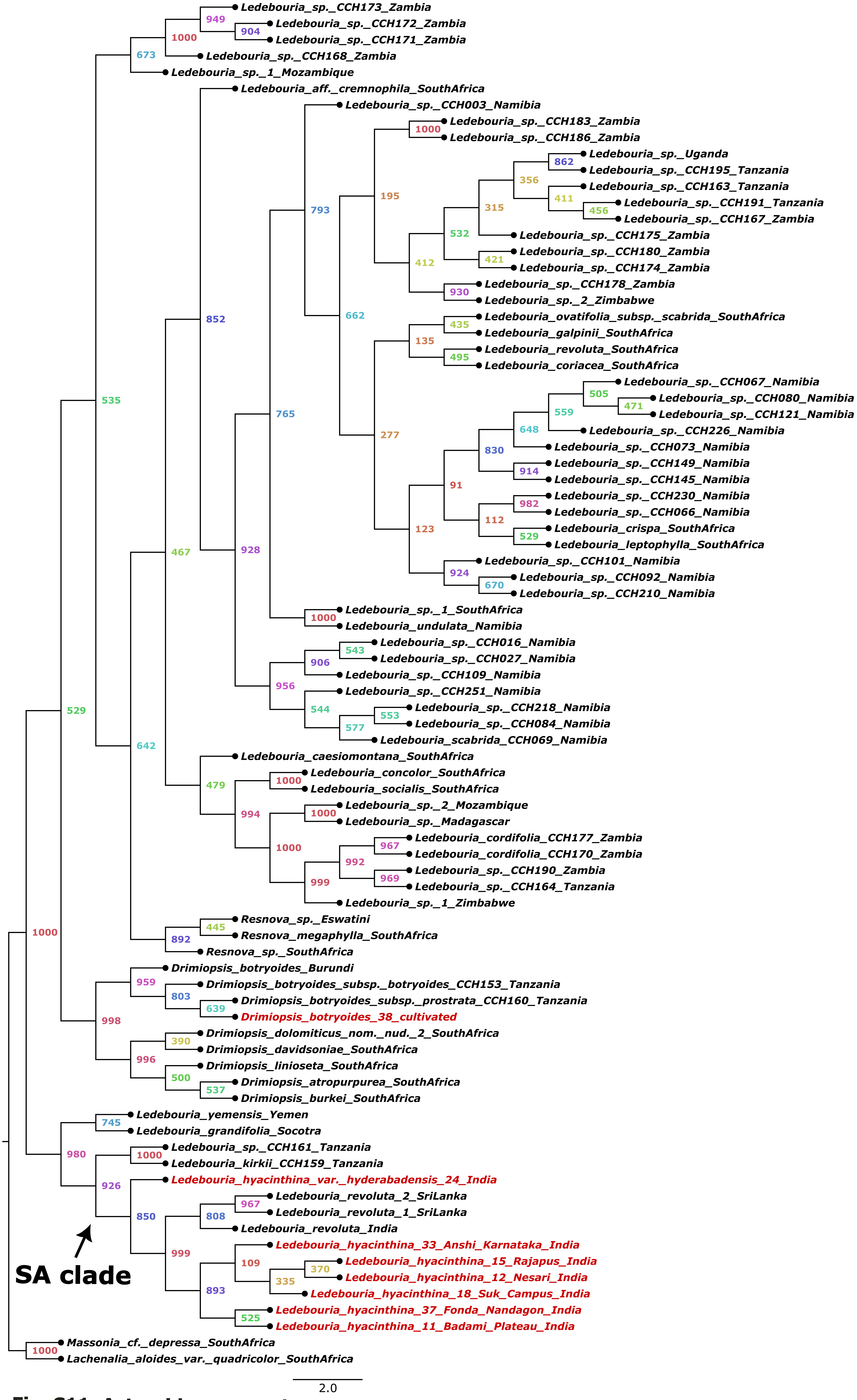


**Fig. S11. Asteroid generated species tree of *Ledebouria* taxa using the untrimmed dataset containing 106 genes. Numbers at nodes indicate bootstrap support. Sequences generated for this study are indicated in red.**

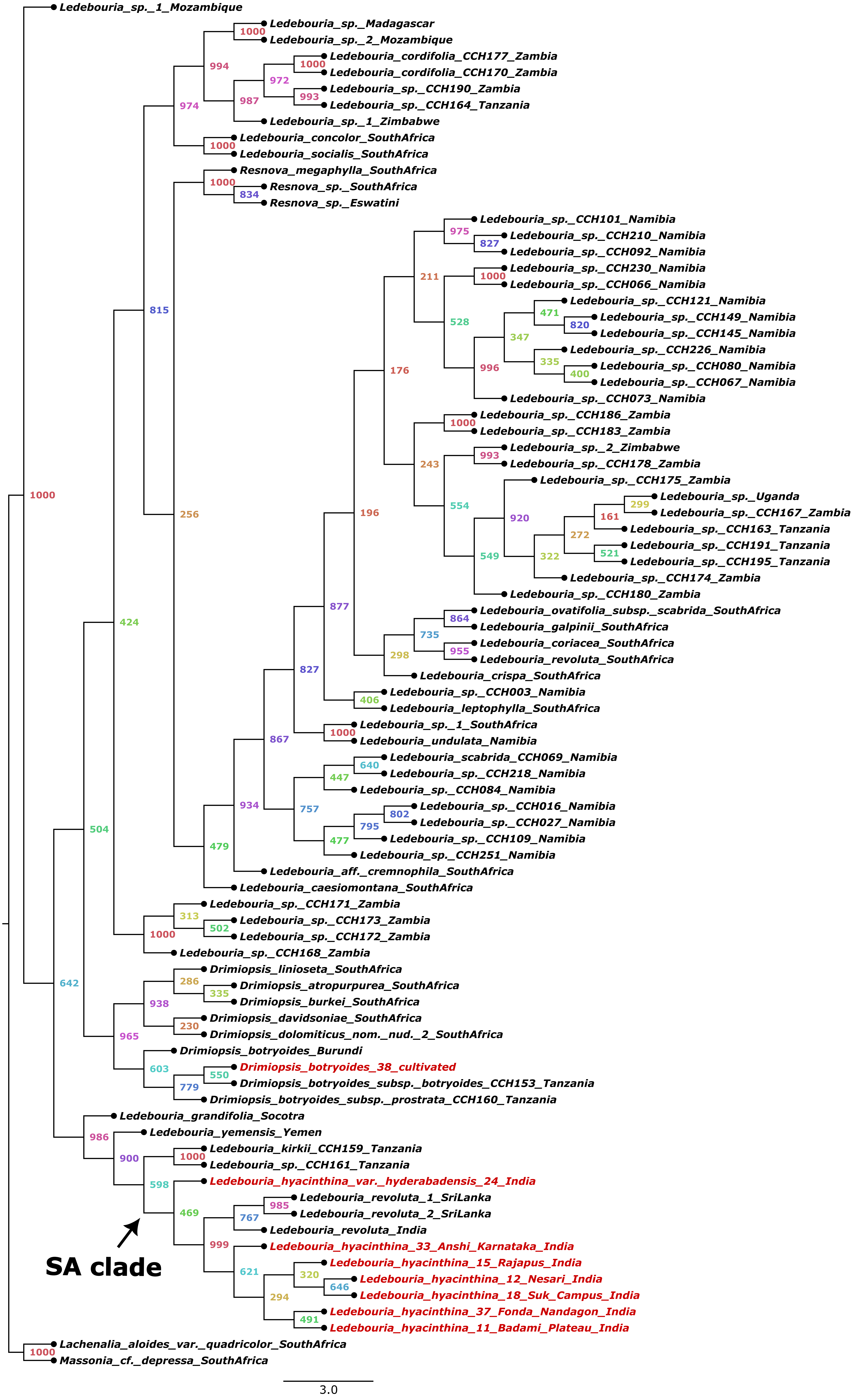


**Fig. S12. Asteroid generated species tree of *Ledebouria* taxa using the trimmed dataset containing 334 genes. Numbers at nodes indicate bootstrap support. Sequences generated for this study are indicated in red.**

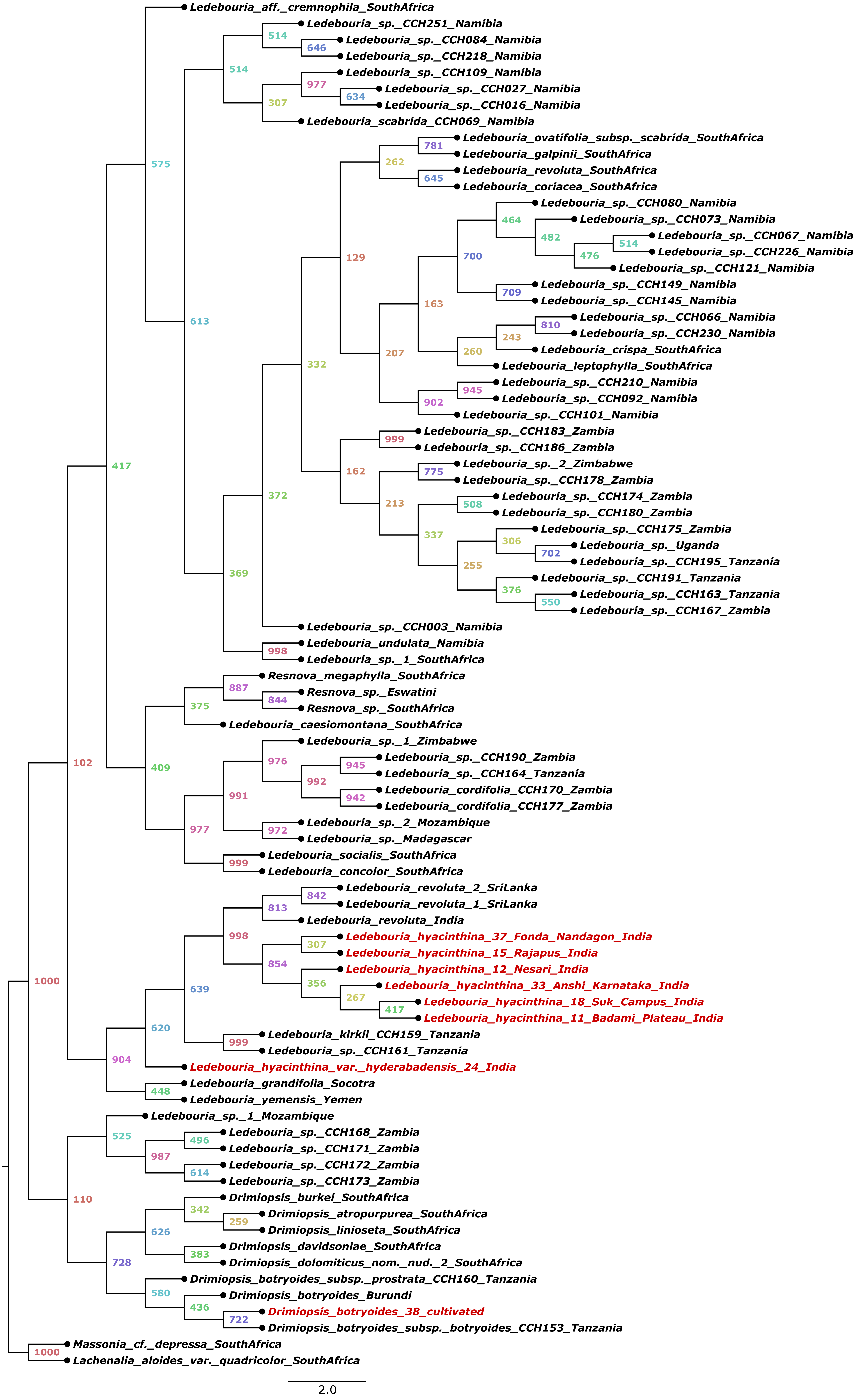


**Fig. S13. Asteroid generated species tree of *Ledebouria* taxa using the trimmed dataset containing 106 genes. Numbers at nodes indicate bootstrap support. Sequences generated for this study are indicated in red.**

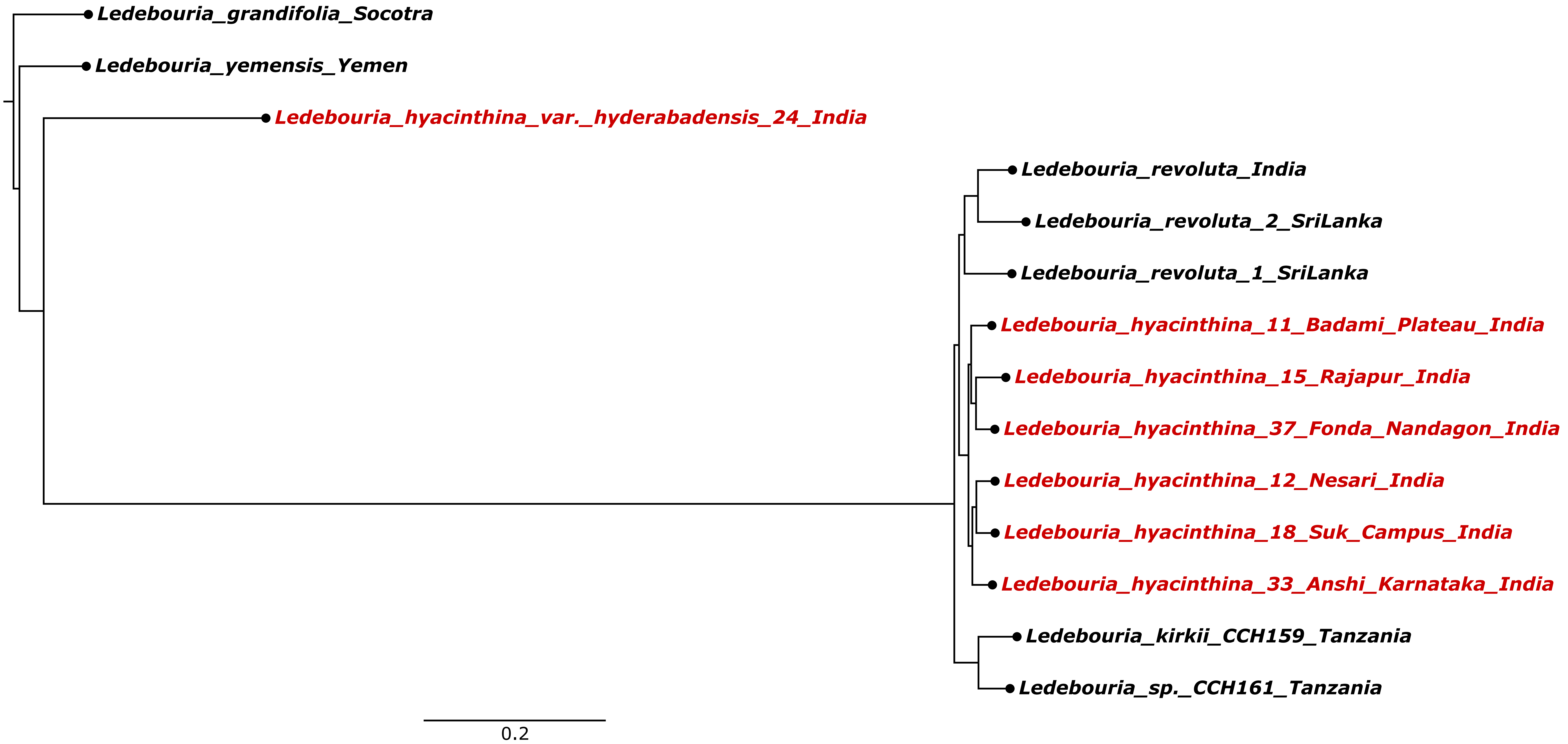


**Fig. S14. NNI-1 species tree for the branch leading to SA clade. Sequences generated for this study are indicated in red.**

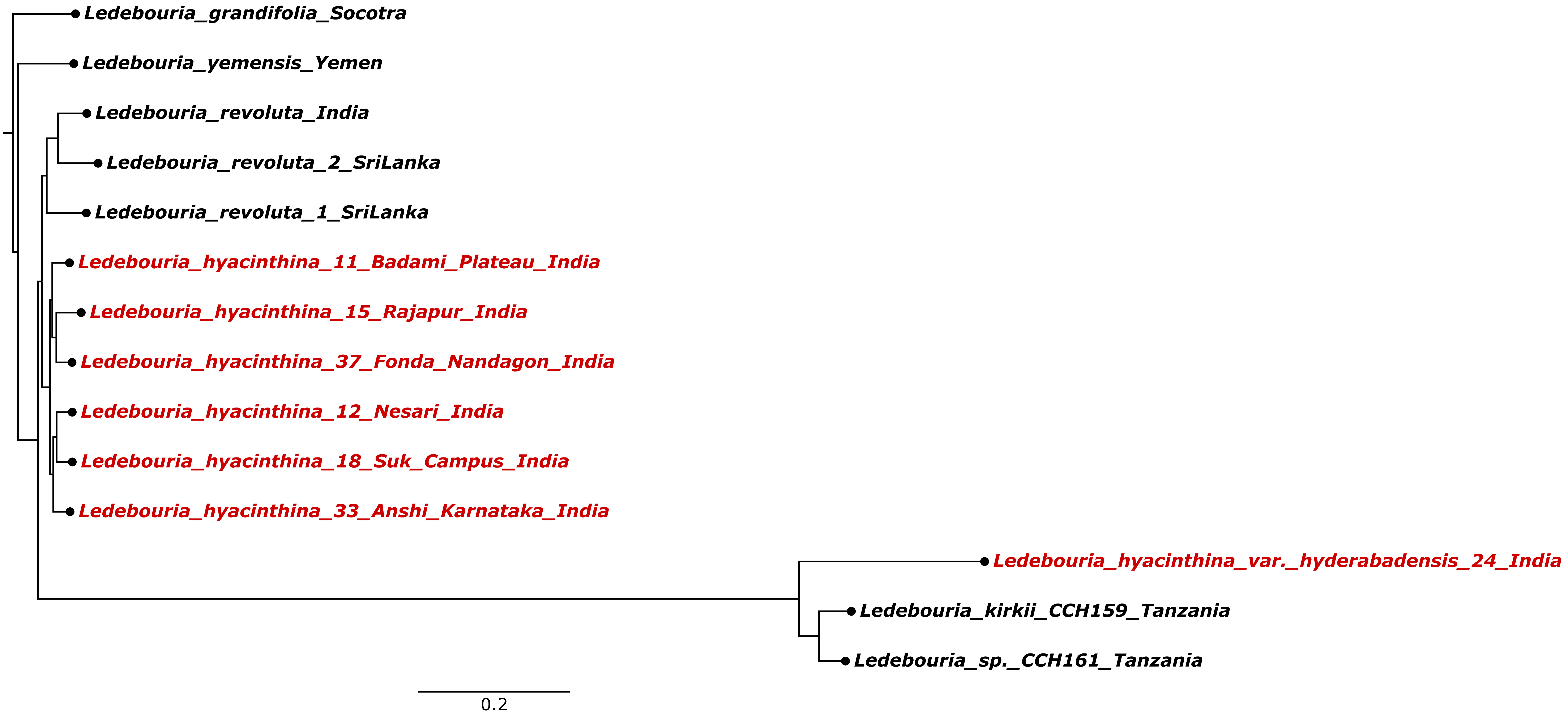


**Fig. S15. NNI-2 species tree for the branch leading to SA clade. Sequences generated for this study are indicated in red.**

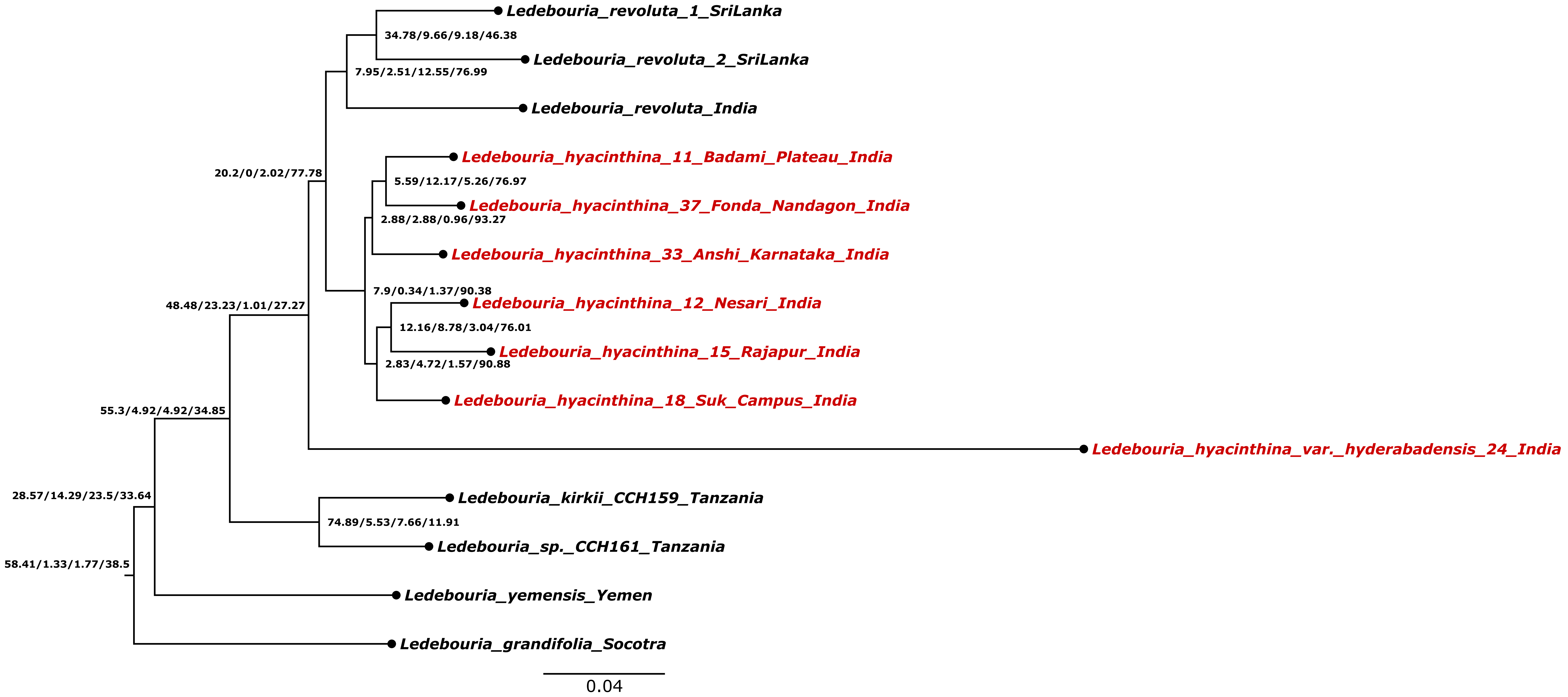


**Fig. S16. Gene concordance factor analysis of the *Ledebouria* SA clade using the untrimmed dataset containing 334 genes. Numbers on the nodes indicate Gene concordance factor/Gene discordance factor for Nearest Neighbour Interchange (NNI)-1 branch/Gene discordance factor for NNI-2 branch/Gene discordance factor due to polyphyly. Sequences generated for this study are indicated in red.**

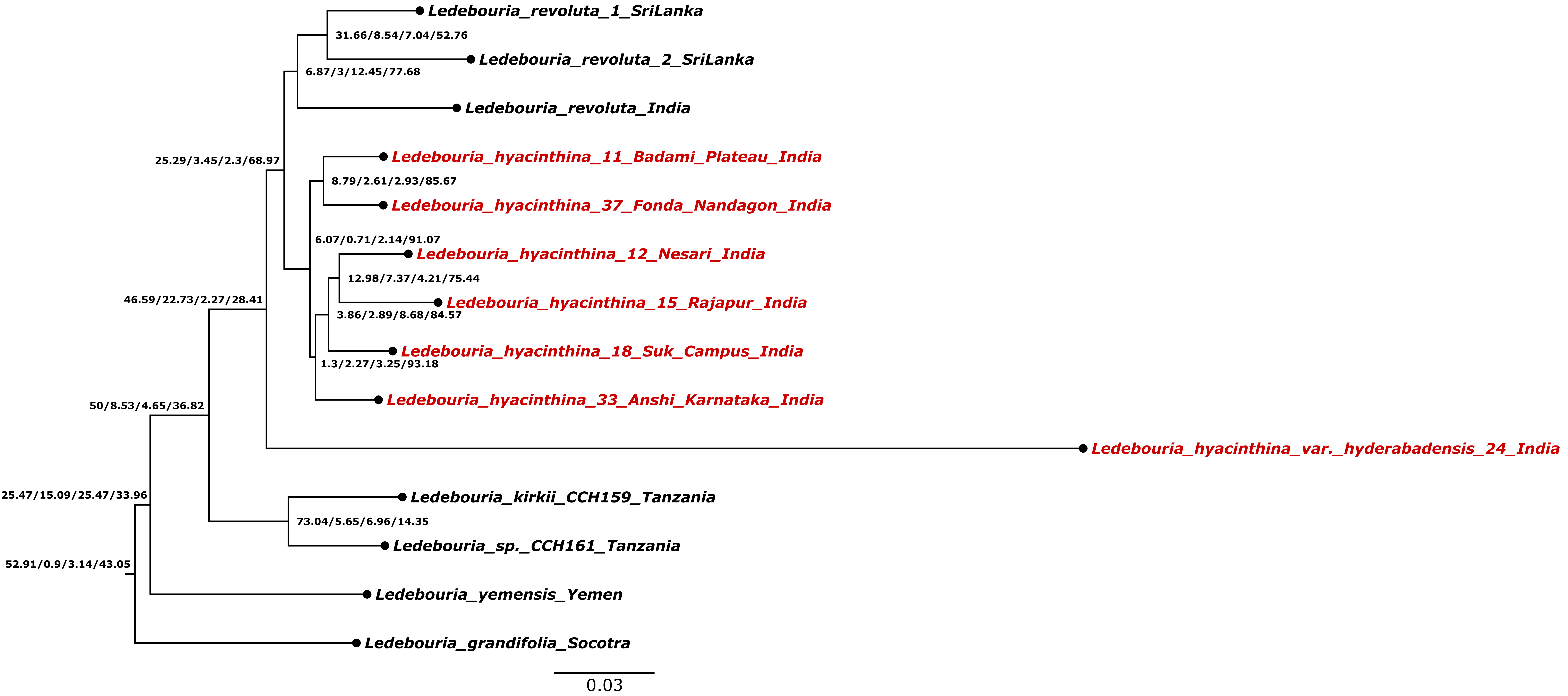


**Fig. S17. Gene concordance factor analysis of the *Ledebouria* SA clade using the trimmed dataset containing 334 genes. Numbers on the nodes indicate Gene concordance factor/Gene discordance factor for Nearest Neighbour Interchange (NNI)-1 branch/Gene discordance factor for NNI-2 branch/Gene discordance factor due to polyphyly. Sequences generated for this study are indicated in red.**

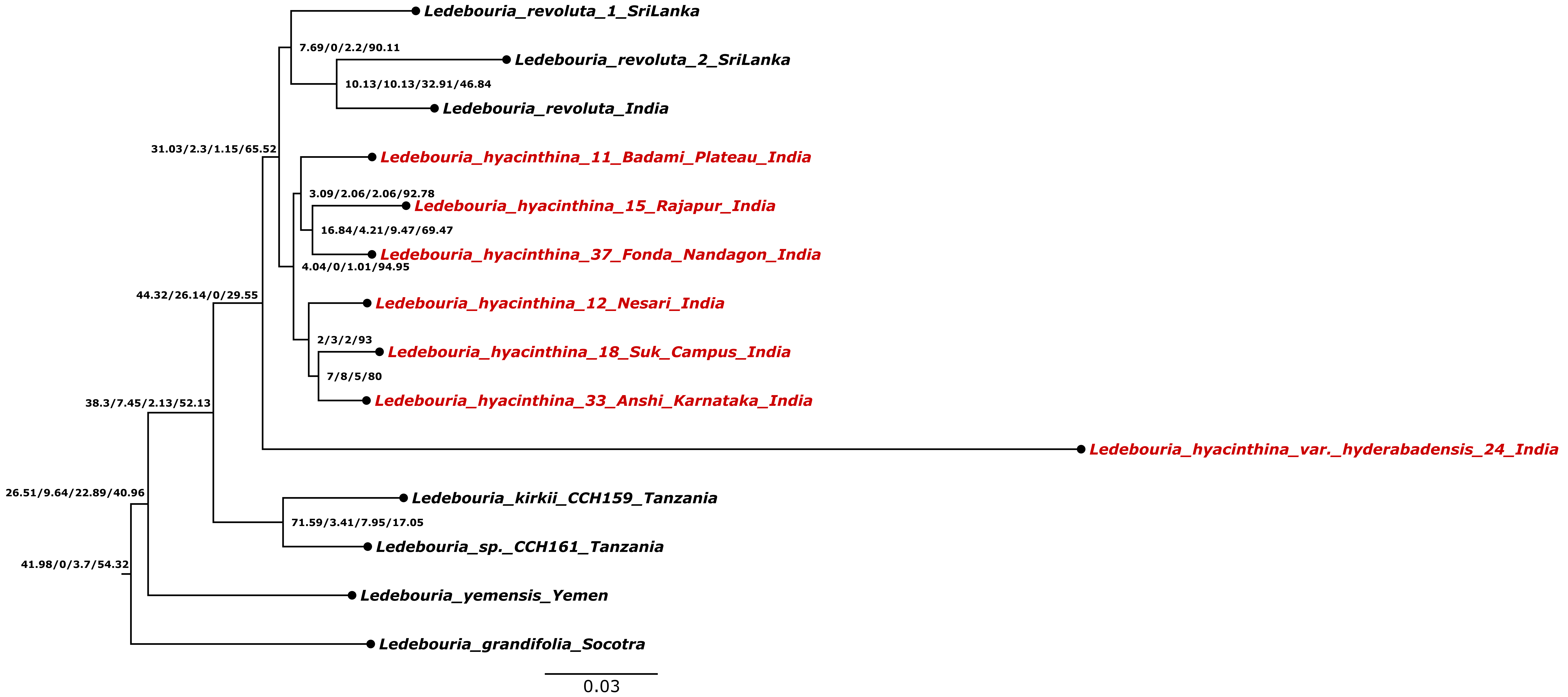


**Fig. S18. Gene concordance factor analysis of the *Ledebouria* SA clade using the trimmed dataset containing 106 genes. Numbers on the nodes indicate Gene concordance factor/Gene discordance factor for Nearest Neighbour Interchange (NNI)-1 branch/Gene discordance factor for NNI-2 branch/Gene discordance factor due to polyphyly. Sequences generated for this study are indicated in red.**

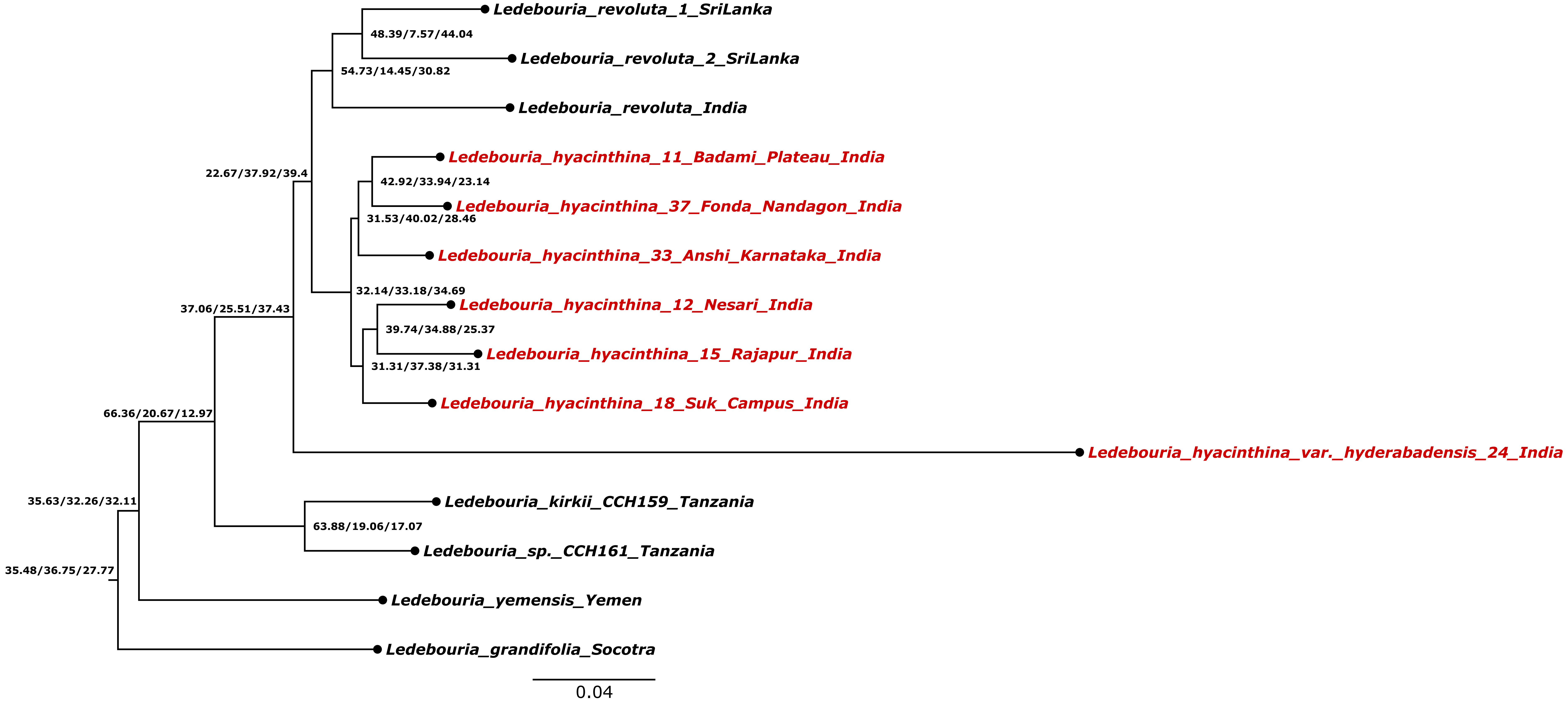


**Fig. S19. Site concordance factor analysis of the *Ledebouria* SA clade using the untrimmed dataset containing 334 genes. Numbers on the nodes indicate Site concordance factor averaged over 100 quartets/Site discordance factor for alternative quartet 1/Site discordance factor for alternative quartet 2. Sequences generated for this study are indicated in red.**

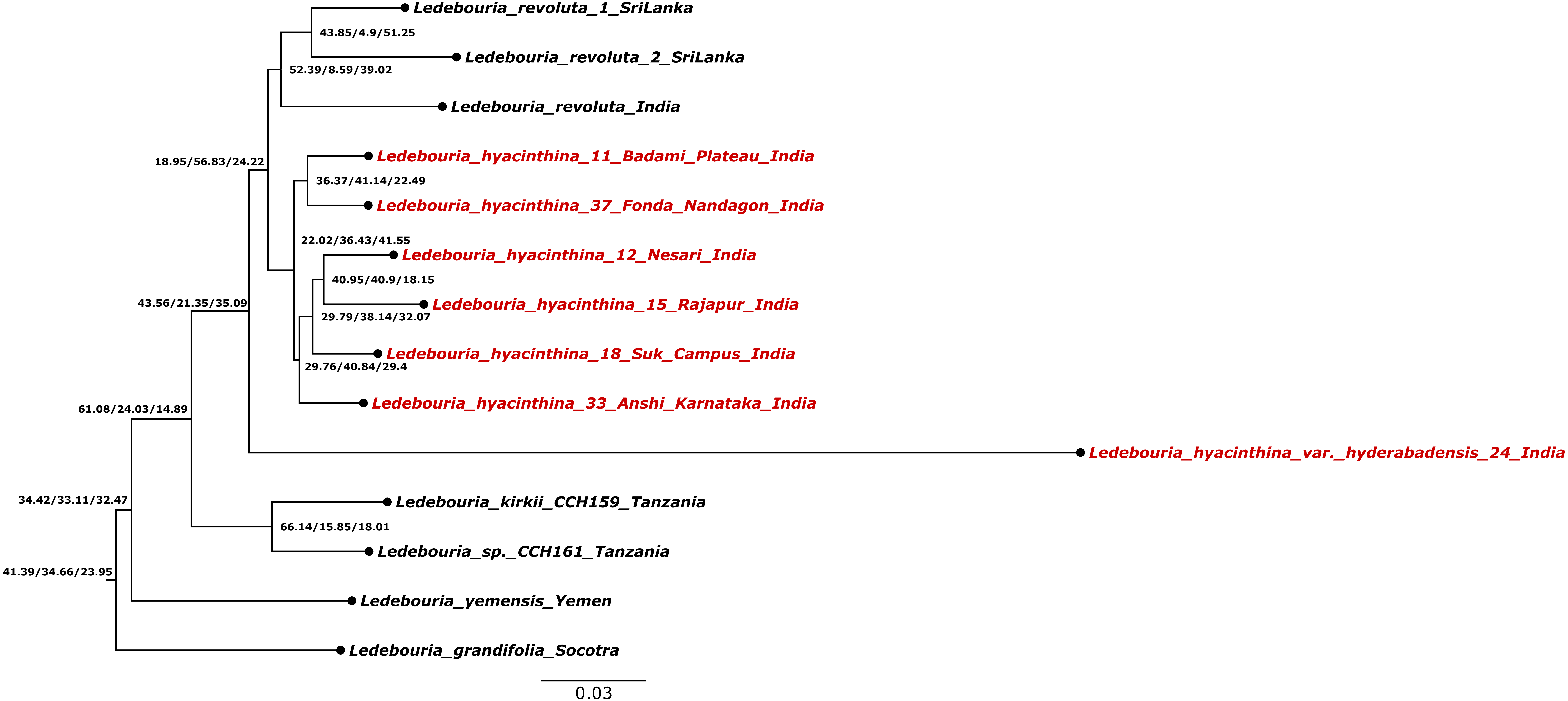


**Fig. S20. Site concordance factor analysis of the *Ledebouria* SA clade using the trimmed dataset containing 334 genes. Numbers on the nodes indicate Site concordance factor averaged over 100 quartets/Site discordance factor for alternative quartet 1/Site discordance factor for alternative quartet 2. Sequences generated for this study are indicated in red.**

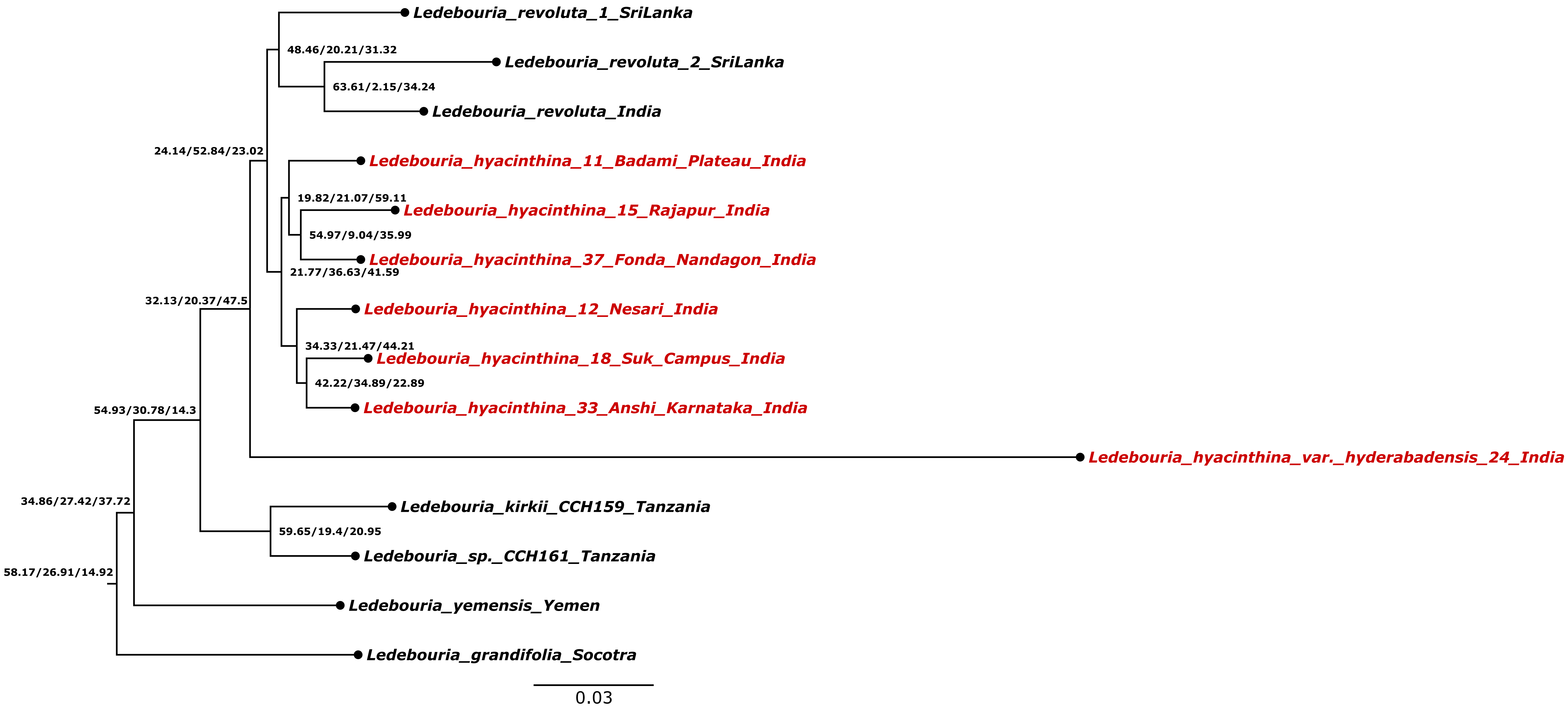


**Fig. S21. Site concordance factor analysis of the *Ledebouria* SA clade using the trimmed dataset containing 106 genes. Numbers on the nodes indicate Site concordance factor averaged over 100 quartets/Site discordance factor for alternative quartet 1/Site discordance factor for alternative quartet 2. Sequences generated for this study are indicated in red.**

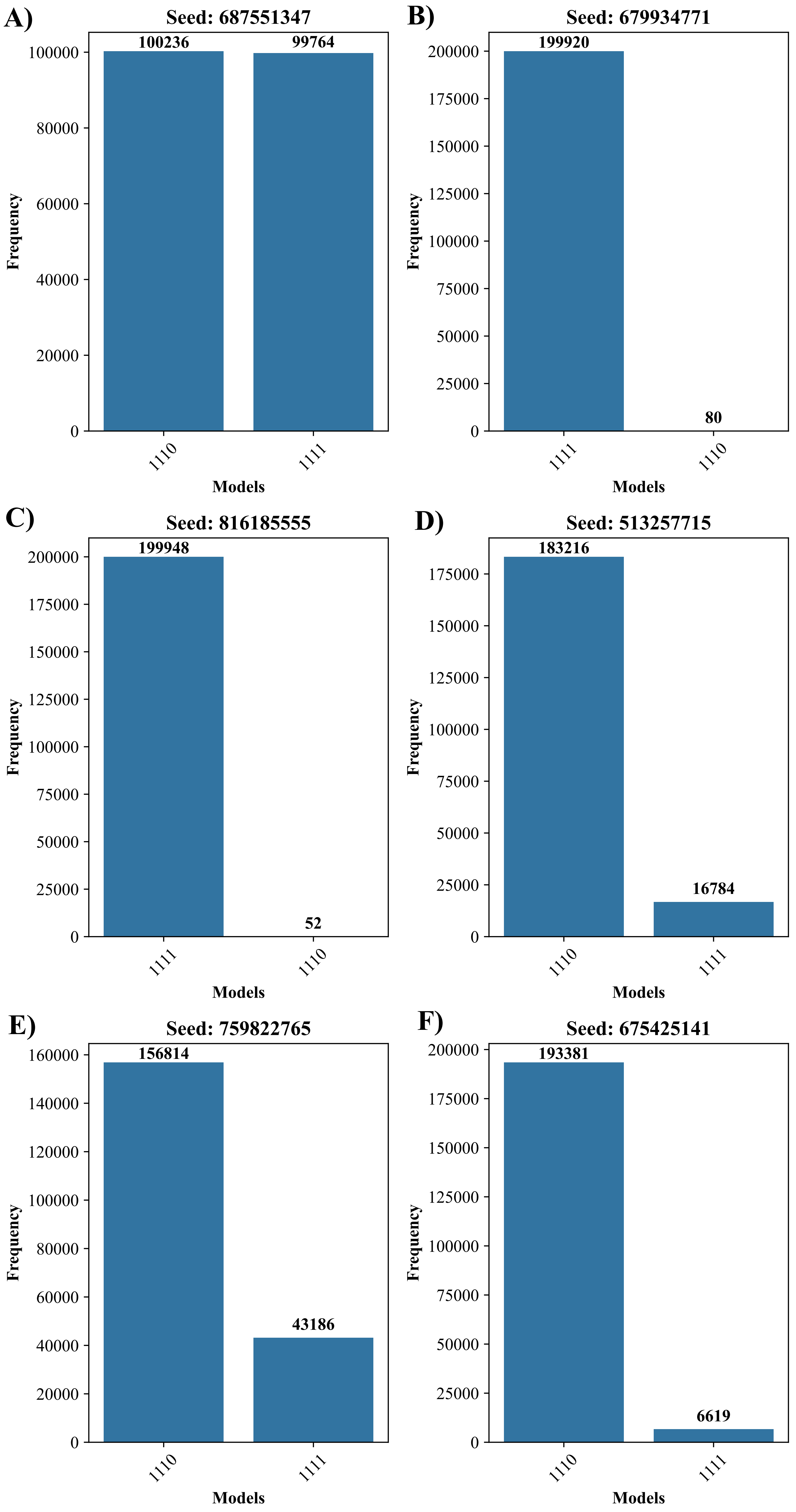


**Fig. S22. Barplots showing the frequency of each model evaluated during species delimitation analysis. (A) and (B): thetaprior=invgamma 3 0.002 int and tauprior=invgamma 3 0.01; (C) and (D): thetaprior=invgamma 3 0.005 int and tauprior=invgamma 3 0.03; (E) and (F): thetaprior=invgamma 3 0.01 int and tauprior=invgamma 3 0.06.**

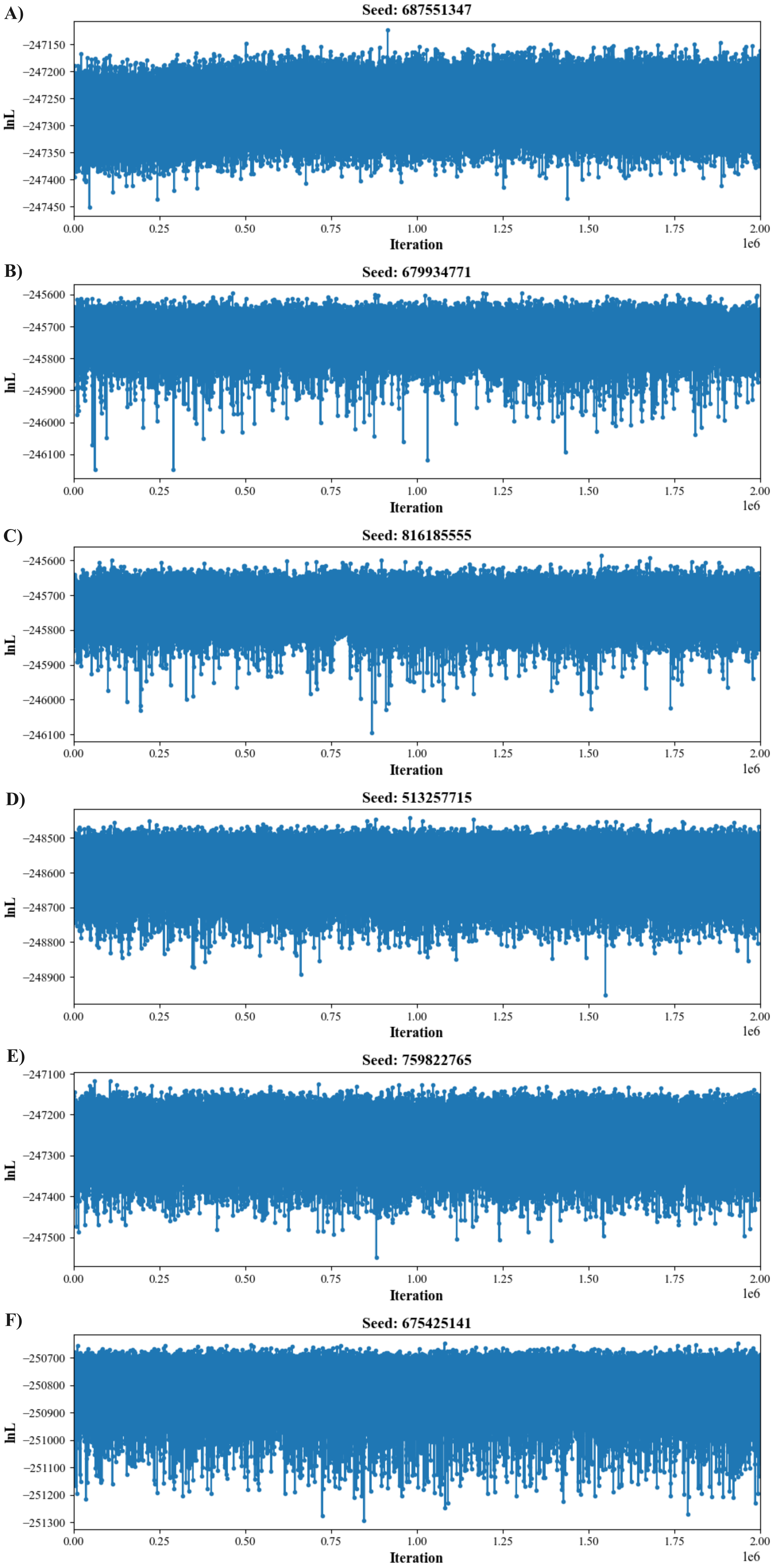


**Fig. S23. Lineplots showing the distribution of lnL across iteration for each model evaluated during species delimitation analysis.**
**(A) and (B): thetaprior=invgamma 3 0.002 int and tauprior=invgamma 3 0.01; (C) and (D): thetaprior=invgamma 3 0.005 int and tauprior=invgamma 3 0.03; (E) and (F): thetaprior=invgamma 3 0.01 int and tauprior=invgamma 3 0.06.**

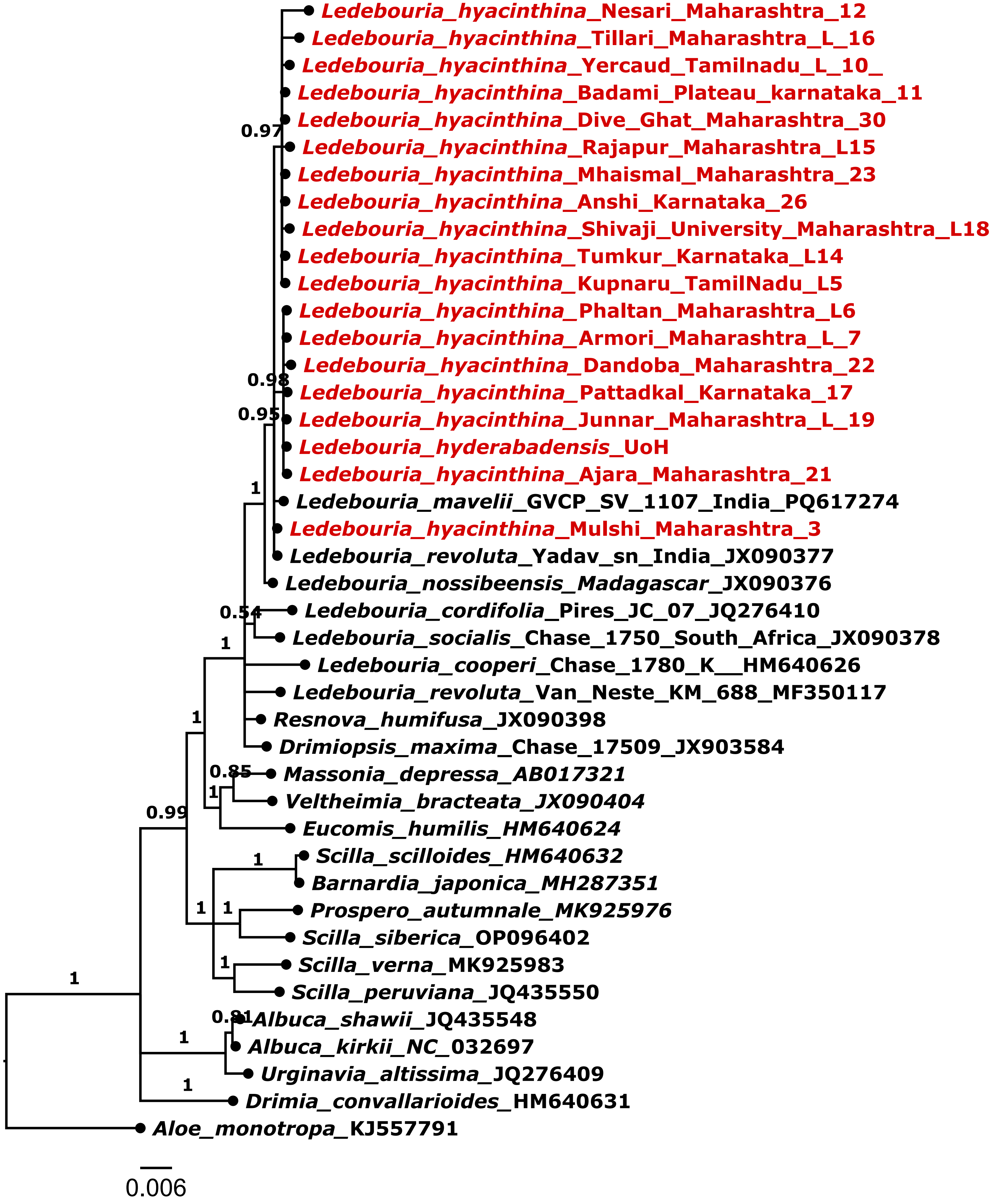


**Fig. S24. Bayesian phylogeny analysis of matK data. Sequences generated for this study are indicated in red.**
